# Characterization of programmed cell death pathways activated in *Mycobacterium tuberculosis*-infected human macrophages

**DOI:** 10.64898/2026.01.30.702894

**Authors:** Guanchao Ding, Jacques Augenstreich, Anushka Poddar, Akshaya Ganesh, Liron David, Ravin Fisher, Volker Briken

**Affiliations:** Department of Cell Biology and Molecular Genetics, University of Maryland, College Park, MD, USA; Department of Life Sciences, Ben-Gurion University of the Negev, Negev, Israel

## Abstract

*Mycobacterium tuberculosis* (Mtb) primarily infects human lung macrophages, which serve as its major replication niche. Mtb can manipulate host macrophage cell death pathways to its advantage by inhibiting apoptosis and inducing necrotic cell death. However, the specific necrotic cell death pathway activated in human macrophages after Mtb infection remains unclear. Here, we used the THP-1 cell line and primary human monocyte-derived macrophage (hMDM) to analyze multiple programmed cell death pathways during days 1-3 after Mtb infection. Confocal microscopic analysis demonstrates that Mtb-infected THP-1 cells or hMDMs rarely exhibited apoptosis. Immunoblotting shows that Mtb induces significant CASP3 and GSDME activation in THP-1 cells, but not in hMDMs. We show that Mtb, in THP-1 cells but not hMDM, induces a significant increase in GSDMD cleavage, a hallmark of pyroptosis. MLKL phosphorylation was not observed in THP-1 cells or hMDMs during Mtb infections, indicating an absence of necroptosis. No changes in ferroptosis markers such as GPX4 expression or lipid peroxidation levels were detected. Time-lapse live-cell imaging revealed no lysosomal membrane permeabilization prior to plasma membrane rupture (PMR). However, we observed DNA release from Mtb-infected THP-1 cells and hMDMs after PMR. The DNA released from THP-1 cells exhibits low levels of myeloperoxidase and histone H3 citrullination. High-resolution confocal imaging shows that Mtb is associated with the released DNA. We demonstrate that pyroptosis induction in THP-1 cells is dispensable for the DNA release and cell death induction. In conclusion, our results reveal that Mtb-triggered cell death in hMDMs bypasses canonical cell death pathways like apoptosis, pyroptosis, necroptosis, and ferroptosis. Instead, cell death in both THP-1 cells and hMDMs correlates with DNA release, potentially through a pathway similar to NETosis in neutrophils.

## Introduction

*Mycobacterium tuberculosis* (Mtb), the causative agent of tuberculosis (TB), was responsible for an estimated 1.25 million deaths in 2023. In the same year, approximately 10.8 million individuals, including 1.3 million children, developed TB [1]. These statistics highlight the resurgence of TB as a leading cause of death from infectious diseases following the COVID-19 pandemic. While traditional anti-mycobacterial therapies have been the cornerstone of TB treatment, they are hindered by several limitations, including prolonged treatment duration (4 – 6 months) and the rise of drug-resistant strains [1,2]. As a result, host-directed therapy (HDT) has emerged as a promising adjunctive approach for treating TB [3–5]. TB pathogenesis and transmission are associated with tissue necrosis [6,7]. Understanding the cellular pathways by which Mtb induces host cell necrosis is fundamental to elucidating the mechanistic basis of tissue necrosis in TB and has informed the development of HDT strategies [8–11].

Macrophages are a diverse and highly plastic population of innate immune cells that play essential roles in host defense against infection, maintenance of tissue homeostasis, and regulation of inflammatory responses [12]. The macrophage heterogeneity affects tuberculosis disease outcomes [13]. Macrophages employ a range of antimicrobial mechanisms, including autophagy, production of reactive oxygen species (ROS) and nitric oxide, acidification of phagolysosomes, and induction of apoptosis, but Mtb can evade elimination by inhibiting and/or resisting these processes [14–16]. Host cell apoptosis is associated with host resistance, but Mtb can inhibit apoptosis induction via several virulence factors [16,17]. For example, the deletion of *nuoG* in Mtb results in increased host cell apoptosis via a ROS/TNF pathway [18]. The *nuoG* deletion mutant has an attenuated phenotype after aerosol infection of mice, supporting the model that host cell apoptosis increases host resistance [18,19]. In contrast, Mtb promotes necrosis, a form of cell death that facilitates bacterial release due to plasma membrane rupture and is associated with host susceptibility [20–22]. After infection, the membrane of the Mtb-containing phagosome can be permeabilized, and in some cases, bacteria escape into the cytosol, and subsequently, host cell necrosis is induced [23,24]. Multiple virulence factors are associated with these processes, such as the type VII secretion systems ESX-1 and ESX-5, the virulence lipid phthiocerol dimycocerosate (PDIM), and the tuberculosis necrotizing toxin [14,16,20]. Nevertheless, the precise host cell signaling pathway by which Mtb induces necrosis is still not fully elucidated.

Necrosis, characterized by the loss of plasma membrane integrity, encompasses a broad spectrum of programmed cell death pathways [25–27]. In macrophages, Mtb induces pyroptosis through the activation of canonical NLRP3 inflammasomes, as demonstrated in human THP-1 cells and murine bone marrow-derived macrophages (BMDMs) [28–32]. In THP-1 cells, this inflammasome activation involves the primate-specific NLRP11 protein and the non-canonical inflammatory caspase-4 and -5 [33]. Conversely, other studies report that Mtb can suppress inflammasome activation via specific virulence factors [34–36]. Mtb may induce necroptosis under certain conditions in BMDMs [37–40]. Increased mitochondrial ROS levels have been associated with gasdermin D (GSDMD)-mediated necroptosis in BMDMs harboring the *Lrrk2^G2019S^* mutation, which is associated with increased risk for Parkinson’s disease in humans [41]. Mtb triggers ferroptosis involving GPX4 and BACH1 pathways in murine *ex vivo* and *in vivo* models, in human monocyte-derived macrophages (hMDMs), and in clinical samples from tuberculosis patients [42–44]. Ferroptosis is associated with tissue damage and increased susceptibility to Mtb [42–44]. In addition, Mtb has been implicated in inducing poorly defined forms of cell death; for instance, lysosomal involvement in Mtb-induced cell death has been observed in TNF-treated human macrophages and untreated murine macrophages [45–47]. The type I interferon (IFN) signaling pathway has been shown to modulate uncharacterized cell death induction during Mtb infection [48,49]. Nuclear deformation and extracellular DNA release have also been reported following Mtb infection, which may suggest the induction of macrophage extracellular trap formation (METosis) via an uncharacterized signaling pathway in human macrophages [47,50,51]. Despite substantial progress in elucidating cell death mechanisms, the specific conditions that determine which death pathways are activated during Mtb infection are still poorly defined. Moreover, the significant variability observed across experimental models contributes to inconsistent findings and complicates interpretations. Notably, two studies comprehensively investigated multiple cell death pathways using BMDMs and chemical inhibitors, yet failed to detect robust activation of any known forms of cell death [49,52], which seemingly contradicts previous reports. Furthermore, many of these studies were performed using murine BMDMs, but Mtb is a human-adapted pathogen, and the host-pathogen interaction regarding cell death signaling might be fundamentally different between murine and human macrophages. For example, type I IFN signaling was involved in cell death induction after infection of murine macrophages but not human macrophages [48,49]. This underscores the need for systematic analyses of the regulation of cell death modalities during Mtb infection in human macrophages.

Our study provides a systematic analysis of the major canonical cell death pathways previously associated with Mtb infection using differentiated THP-1 cells and primary hMDMs. In THP-1 cells, Mtb infection significantly induces apoptosis and pyroptosis. In contrast, these pathways are not activated in hMDMs at the examined time points post-infection. Neither necroptosis nor ferroptosis was detected in either cell type over a period of 3 days post-infection. Live-cell confocal microscopy was performed to image the lysosomal pH and track single Mtb-infected cells during the final two hours preceding plasma membrane rupture (PMR), and no detectable lysosomal membrane permeabilization (LMP) was observed. However, this analysis revealed DNA release events associated with necrotic cell death. Subsequent experiments confirmed the presence of DNA release in both THP-1 cells and hMDMs during Mtb infection. In THP-1 cells, this phenomenon accounted for around 60% of overall necrotic cell death. In contrast, in Mtb-infected hMDMs, only about 30% of cells exhibited DNA release, suggesting that the dominant form of early necrotic death in hMDMs remains mechanistically uncharacterized. Our findings further indicate that the released DNA can bind to extracellular Mtb. Additionally, we show that THP-1 cells deficient in caspase-4 and -5 (*CASP4/5*) or ninjurin-1 (*NINJ1*) exhibit DNA release and cell death following Mtb infection to an extent comparable to that observed in wild-type THP-1 cells. In conclusion, our study identifies a correlation between DNA release and necrotic cell death modality in THP-1 and hMDM cells during Mtb infection. The precise signaling pathways remains to be characterized but we show that pyroptosis, necroptosis or ferroptosis necrosis signaling pathways are not involved.

## Materials and Methods

### Cell cultures and differentiation

Wild-type (w.t.) THP-1 cells (TIB-202) were obtained from ATCC (American Type Culture Collection; VA, USA). THP-1 *ninj1*^−/−^ mutant cells were generated as described [53]. The THP-1 *casp4/5*^−/−^ strain and its corresponding w.t. control were kind gifts from Dr. Clare Bryant (University of Cambridge, UK) [54]. Monocytic THP-1 cells were cultured in RPMI 1640 medium (ATCC, 30-2001) supplemented with 10% heat-inactivated FBS (Gibco, NY, USA; A52567-01). The culture medium was refreshed every 2–3 days to maintain a cell density between 2×10^5^ and 1×10^6^ cells/mL prior to differentiation. THP-1 cells were pelleted and resuspended in complete medium containing 50 ng/mL phorbol 12-myristate 13-acetate (PMA) (Sigma, MA, USA; P1585), adjusted to a final density of 1×10^6^ cells/mL. Cells were seeded at 5×10^5^/well in non-treated 24-well plates (ThermoFisher, MD, USA; 144530) or 2×10^5^/well in uncoated imaging chambers (ibidi, Gräfelfing, DE; 80821). Cells were treated with PMA for 2 days, after which they were washed twice with Dulbecco’s phosphate-buffered saline (DPBS; ThermoFisher, 14190144) and incubated for an additional 2 days in fresh PMA-free medium before treatment or infection. All cell lines were stored in freezing medium composed of 10% dimethyl sulfoxide (DMSO) in 90% FBS for cryopreservation.

Human peripheral blood mononuclear cells (PBMCs) were isolated from LeukoPak according to the protocol as previously described [55]. CD14+ monocytes were purified using CD14 MicroBeads (Miltenyi Biotech, CA, USA; 130-050-201), with the bead volume reduced to one-third of the manufacturer’s recommended amount. All other steps were performed according to Miltenyi Biotech’s instructions. To obtain human monocyte-derived macrophages (hMDMs), monocytes were pelleted and resuspended at 1×10^6^ cells/mL in hMDM growth medium consisting of RPMI 1640, 10% FBS, 5% human AB serum (GeminiBio, CA, USA; 100-318-100), and 10 ng/mL human M-CSF (Bio-Techne, MN, USA; 216-MCC-010). Monocytes were seeded in uncoated 24-well plates or uncoated imaging chambers at the same density as THP-1 cells. On day 3 post-seeding, additional growth medium was added, and the medium was subsequently replaced every 3 days. hMDMs were used for the following experiments on day 7 or 9 post-differentiation.

Human neutrophils were isolated from whole blood using the EasySep™ Direct Human Neutrophil Isolation Kit (Stemcell Technologies, Van, CA; 19666). After isolation, neutrophils were washed once with THP-1 growth medium to remove residual EDTA and seeded into 8-well collagen I-coated imaging chambers (ibidi, 80829). Cells were seeded for at least 1 hour to allow adherence before Mtb infection. Refer to Appendix Table 2 for detailed information on materials, kits, and reagents.

The HT-29 cell line (ATCC, HTB-38; LOT: 70061450) was cultured in the same growth medium as THP-1 cells. Accutase (Millipore-Sigma, A6964) was used at 10 mL per 75 cm² flask to detach HT-29 cells for passaging or seeding. Cells were seeded at the same density as THP-1 cells 1 d prior to necroptosis induction. HT-29 cells were passaged every 3–4 d upon reaching approximately 80% confluence.

### Cell treatments and experimental controls

THP-1 cells were treated with human recombinant IFN-β (R&D, 8499-IF-010) at the indicated concentrations and times. Raptinal (Sigma-Aldrich, SML1745) was administered at 10 μM for 4–6 h to induce cleavage of CASP-3, CASP-9, and GSDME. Staurosporine (Sigma-Aldrich, S6942) was administered at 2 μM for 6 hours to induce cleavage of CASP-3, CASP-8, and CASP-9. To induce GSDMD cleavage and pyroptosis, THP-1 cells were treated with LPS (InvivoGen, CA, USA; tlrl-eblps) at 100 ng/mL for 4 h, treatment with 20 μM nigericin (InvivoGen, tlrl-nig) for 30 min. To inhibit GSDMD activation, THP-1 cells were treated with disulfiram (MedChemExpress, HY-B0240) at 20 μM. For necroptosis induction, THP-1 cells were pretreated with z-VAD-FMK (Cell Signaling Technology, 60332S) at 20 μM for 1 h, followed by treatment with 100 ng/mL human TNF-α (R&D Systems, 10291-TA-020) in combination with 10 μM SM-164 (Cell signaling Technology, 56003S). For ferroptosis induction, THP-1 cells were treated with 10 μM RSL-3 (MedChemExpress, HY-100218A). Where indicated, ferroptosis was inhibited by the addition of 10 μM ferrostatin-1 (MedChemExpress, HY-100579). For LMP induction, THP-1 cells were treated with LLoMe (BACHEM, 4000725) at 1 μM for at least 1 hour. All treatments were conducted at 37 °C in a humidified atmosphere with 5% CO₂.

### Mycobacterial strains and cultures

*M. tuberculosis* H37Rv, *M. smegmatis* mc²155, and *M. kansasii* (ATCC, 12478) were cultured in Middlebrook 7H9 broth (BD, CT, USA; 271310) supplemented with 0.2% glycerol (Fisher, G33-4), 10% OADC (BD, 21235), and 0.05% Tween-80 (Fisher, MO, USA; BP338-500). More than 5 mL of bacterial culture was transferred to a 50 mL conical tube and incubated at 37 °C with shaking at 140 rpm. Cultures were terminated after four passages. For dsRed-expressing H37Rv, 100 μg/mL of Zeocin (InvivoGen, ant-zn) was added to maintain plasmid selection.

### Infection experiments

Bacterial cultures were centrifuged at 2 000 × g for 7 minutes and resuspended in 0.05% Tween-80 in DPBS (PBS-T). The suspension was subsequently centrifuged at 80 × g for 3 minutes to remove large aggregates. Optical density (OD) was measured using a spectrophotometer, with bacterial concentration estimated based on the assumption that an OD of 1 corresponds to 3 × 10⁸ bacteria/mL.

On the day of infection, culture media were removed, and cells were washed twice with DPBS. THP-1 cells or hMDMs were then infected with Mtb in infection medium at an MOI of 5:1 or 2:1. Infected macrophages were incubated at 37 °C in a humidified atmosphere containing 5% CO₂.

For pulse-chase infections, cells were allowed a 4-hour phagocytosis window, after which they were gently washed twice with 1× PBS and then incubated with chase medium. The chase medium consisted of the growth media supplemented with 100 μg/mL Gentamicin (Corning, NY, USA; 30-005-CR). The 0-hour post-infection (hpi) time point was defined as the end of the 4-hour phagocytosis period. In contrast, for continuous infections, the 0-hour hpi time point was defined as when bacteria were first added to the cells. The infection medium for THP-1 cells consisted of RPMI 1640 supplemented with 10% heat-inactivated FBS and 5% human AB serum (GeminiBio, 100-512-100). For hMDMs, the infection medium was identical to their growth medium.

Lysosomal membrane permeabilization (LMP) induced by Mtb was assessed using LysoView™ 633 (LV; Biotium, CA, USA; 70058) in combination with time-lapse live-cell imaging. LV (concentration unknown) was reconstituted in 100 μL dH₂O following the manufacturer’s instructions and used at a 1:2000 dilution. SYTOX™ Green Nucleic Acid Stain (Syt G; Invitrogen; S7020) was added at 0.5 μM (1:10,000 dilution) to detect plasma membrane rupture.

### LDH and AK release assays

Cell membrane rupture was assessed using the LDH assay kit (Promega, WI, USA; G1780) and the AK release assay kit (Lonza, MD, USA; LT07-217), following the manufacturers’ instructions. Cell culture supernatants were pooled before aliquoting for the respective assays, and 3 parallel wells were set for each experimental condition. Chase medium was used to determine the background absorbance or luminescence intensity. For the LDH release assay, absorbance at 490 nm was measured, while luminescence was recorded for the AK release assay, both using a BioTek plate reader.

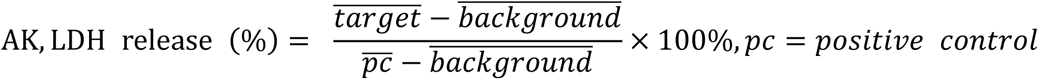

### IFN-β neutralization assay

Following optimization, the IFNAR-neutralizing antibody IFNAR2 Monoclonal Antibody MMHAR-2 (Invitrogen, 213851) was used at 100 ng/mL to block IFNAR signaling. For the isotype control condition, 100 ng/mL of Mouse IgG2a kappa Isotype Control (eBM2a) (Invitrogen, 14-4724-82) was added to THP-1 cells. Antibodies were preincubated with THP-1 cells for 1 hour prior to infection.

### ELISA

The Human IFN-β DuoSet ELISA kit (Bio-techne, DY814-05) and the Human IL-1β DuoSet ELISA kit (Bio-techne, DY201) were used to quantify the concentrations of secreted IFN-β and IL-1β from THP-1 cell supernatants, following the manufacturer’s protocols. Supernatants were pooled and aliquoted for analysis, with each condition assessed in four parallel wells. Absorbance was measured at 450 and 540 nm (background), and final cytokine concentrations were calculated using linear regression in GraphPad Prism.

### YO-PRO-1 and DRAQ7 staining and analysis

THP-1 cells or hMDMs were seeded in 8-well chamber slides and infected with dsRed H37Rv *Mtb* at an MOI of 5 for pulse-chase experiments or left uninfected. At the indicated time points, the culture medium was replaced with 200 μL of staining solution containing 0.5 μM YO-PRO-1 (Biotium) and 0.3 μM DRAQ7™ (Invitrogen, D15106). Cells were stained at 37 °C for 15 min and immediately imaged. For each condition and experimental repeat, more than 200 cells were analyzed.

### SDS-PAGE and Western Blotting

THP-1 cells or hMDMs were lysed in 1% NP-40 lysis buffer (1% NP-40, 0.4 mM EDTA, 10 mM Tris-HCl, 150 mM NaCl) supplemented with Halt™ Protease and Phosphatase Inhibitor Cocktail (Thermo Scientific, 78442). Lysates were centrifuged, and the supernatants were collected. Cell lysates were mixed with 4× Laemmli buffer (Thermo Scientific, J60015.AD) and denatured for 30 minutes before gel-loading.

Total protein concentrations were determined using the Pierce™ BCA Protein Assay Kit (Thermo Scientific, 23227). For SDS-PAGE, 20 µg of protein per sample was loaded onto SurePAGE Bis-Tris 12% gels (GenScript, M00668). Electrophoresis was performed at 50 V for 20 minutes, followed by 150 V for 50 minutes. Proteins were transferred to PVDF membranes (Thermo Scientific, 88585) using the eBlot™ L1 transfer system (GenScript, L00686).

Membranes were blocked with SuperBlock™ Blocking Buffer (Thermo Scientific, 37580) for 20 minutes and incubated with primary antibodies diluted in TBS-T (2.4 g/L Tris-base, 8.8 g/L NaCl, 0.1% Tween-20; pH 7.6) for 2 hours or overnight. After six 2-minute washes with TBS-T, membranes were incubated with HRP-conjugated secondary antibodies in TBS-T for 1 hour, followed by another series of six 2-minute washes. Signal detection was carried out using SuperSignal™ West Pico PLUS Chemiluminescent Substrate (Thermo Scientific, 34580), and membranes were imaged with the iBright™ FL1500 Imaging System (Invitrogen, A44241). The information on antibodies is provided in the Appendix Table 1.

Band intensities were quantified using ImageLab software. Target protein intensities were normalized to β-actin levels from the same experiment, and these values were further normalized to the untreated or uninfected day 1 control for statistical analysis.

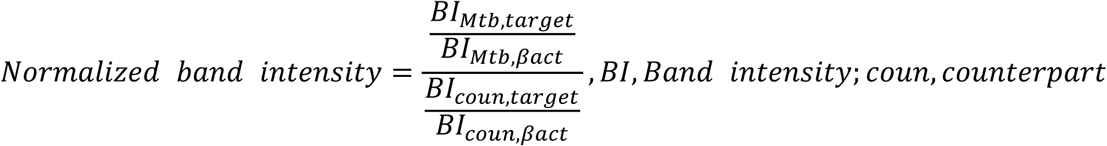

### Liperfluo Assay

Liperfluo (Dojindo Molecular Technologies, MD, USA; L248-10) was added to THP-1 cells at a final concentration of 10 μM, 30 minutes prior to infection. Following incubation, cells were washed twice with DPBS to remove excess dye. DRAQ7™ (Invitrogen, D15106) was added to the infection medium at a 1:1000 dilution to monitor cell death. As a positive control for lipid peroxidation, 100 μM cumyl hydroperoxide (Thermo Scientific, T06I072) was added to uninfected cells 2 hours before imaging.

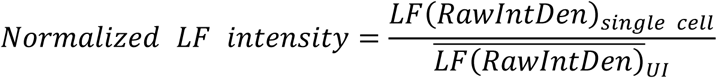

### Image analysis

For infection experiments, time 0 was defined as the frame in which a cell first became positive for Syt G. LV fluorescence intensities were quantified across 12 frames preceding this event for each cell. For control experiments, time 0 corresponded to the first frame of image acquisition. Single-cell segmentation and tracking were conducted using the method and code adapted from “da_tracker.” [56] FIJI and TrackMate-Cellpose combo were combined to quantify the fluorescent intensities.

Cells were classified as “infected” if z-projection images consistently showed dsRed-labeled Mtb within the same cell across multiple consecutive frames. For uninfected cells, 12 consecutive data points were randomly selected from each trackable single cell to establish a baseline. Curves showing more than four data points below the baseline, with statistically significant deviation, were considered indicative of LMP.

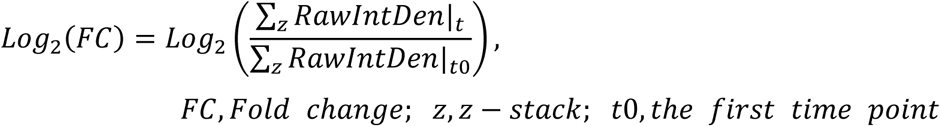

ZEN 3.6 software was used for 3D volume rendering, image generation, and video creation. For 3D visualization, the rendered model was rotated 360° along the Z-axis (as defined by the microscope), with one frame captured per degree of rotation.

Cells exhibiting DNA release were distinguished from cells without DNA release based on the z-projection of the Syt G signal. A “sum” projection was generated from the original 11 z-slices using the Z-Project function in Fiji. If the Syt G signal from an analyzed cell extended beyond the cellular outline, the cell was classified as exhibiting DNA release. For clarity of data presentation, only the z-projection generated using the “max” method is shown, rather than the “sum” projection.

### Immunofluorescence microscopy

Cells cultured in 8-well chambers were first washed once with phosphate-buffered saline (PBS), then fixed with 4% paraformaldehyde (PFA) in PBS (Thermo Scientific, J61899.AK) for 1 hour at room temperature (RT) or overnight at 4°C. After fixation, cells were washed three times with 0.1 M glycine in PBS (PBS-G), with each wash lasting 5 minutes. To block Fc receptors, Human TruStain FcX™ (BioLegend, CA, USA; 422302) was diluted 1:1000 in 3% BSA, 0.2% Tween-20 in PBS (PBS-BT) and applied to the cells for 1 h. Primary antibodies were diluted in 1% BSA in PBS (PBS-B) and incubated with the cells for 2 hours at RT or overnight at 4°C. After three washes with PBS, secondary antibodies and 1 μg/mL 4′,6-diamidino-2-phenylindole (DAPI; Thermo Scientific; 62248), also diluted in PBS-B, were added. Cells were washed three additional times with PBS after the staining. Information on the antibodies used is listed in Appendix Table 1. The DAPI and T-PMT channels were used as references to manually define regions of interest (ROIs) for mean fluorescence intensity (MFI) quantification in FIJI.

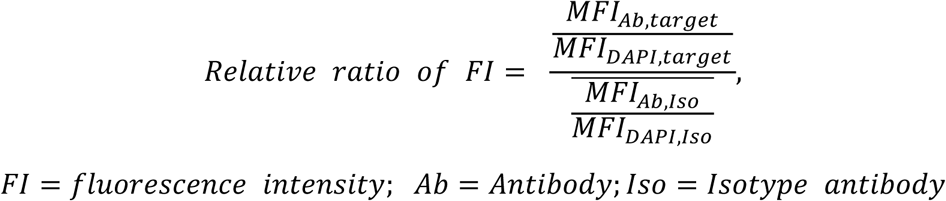

### Confocal microscopy

Cells were imaged using a Zeiss LSM 800 laser scanning confocal microscope equipped with two gallium arsenide phosphide photomultiplier tube (GaAsP-PMT) detectors and one transmitted light photomultiplier tube (T-PMT) detector. A 40×/NA 1.2 water immersion objective was used for all imaging experiments.

For time-lapse live-cell imaging, the observation chamber was maintained at 37°C with 5% CO₂ supplied via a humidified incubator. Z-stacks spanning 10 μm with 1 μm intervals were acquired every 10 minutes for up to 20 hours. Each field consisted of 11 optical slices. A total of four fields were imaged, with a maximum of two experimental conditions per experiment. Each image spanned an area of 319.45 × 319.45 μm, corresponding to 2048 × 2048 pixels. For immunofluorescence microscopy, 3-4 fields were imaged with Z-stacks spanning 20 μm with 1 μm interval. 1 slice with proper signals was selected for MFI quantification. Airyscan confocal microscopy and 3D volume rendering were performed using a Zeiss LSM 980 microscope with the same 40×/NA 1.2 water objective.

### CFU analysis

THP-1 cells were infected with w.t. H37Rv Mtb at an MOI of 5 for 4 h, after which the infection medium was replaced with fresh growth medium without gentamycin. DNase I (100 U/mL) was added to the wells either immediately after infection (0 h post-infection) or 1 h prior to supernatant collection at 2 or 3 d post-infection. Supernatants were diluted in DPBS at 1:10 and 1:100, and 100 μL of each dilution was plated in triplicate on 7H11 agar plates. Plates were monitored daily to ensure the absence of contamination, and CFUs were enumerated at 24 d post-inoculation. Only plates containing more than 20 countable colonies were included in the analysis. For each experimental condition, the mean CFU value of all technical replicates was used as the representative CFU.

### Statistics

All statistical analyses were performed using the built-in analysis modules in GraphPad Prism (version 10.0 or later), except for the modified Chi-square analysis. The modified Chi-square test was used to compare two curves with known or assumed means and standard deviations (SDs), and was performed either with the Excel spreadsheet provided in a previous publication [57] or automated by Python programming for batch analyses. In this study, each independent experiment (for THP-1 and HT-29) or donor (for hMDM) represents a biological replicate. Grubb’s test at α = 0.05 was applied to identify at most 1 outlier for WB analysis. For experiments involving hMDMs, samples from at least three donors were used. In bar graphs, bar height indicates the mean value of the data. A *p*-value < 0.05 was considered statistically significant.

### Data and Code availability

The videos and image analysis files are available at https://data.mendeley.com/, DOI: 10.17632/rkbw676zf2.3

## Results

### Inhibition of Interferon-α/β receptor (IFNAR) signaling does not prevent Mtb-induced cell death in THP-1 cells

In order to systematically investigate the activation of several established cell death pathways after Mtb infection in human macrophages, we first performed a kinetic analysis of host cell death induction (Fig.1A, B). Adenylate kinase (AK) and lactate dehydrogenase (LDH) release assays were performed as readouts for cell death following infection with Mtb at a multiplicity of infection (MOI) of 5 for the indicated durations (Fig. 1A, B). We noticed a significant increase in cell death after 1 day of infection in Mtb-infected versus uninfected THP-1 cells and hMDM that peaked around day 3. No further increase in AK or LDH release in the Mtb group was observed at days 4 or 5 compared with day 3, likely due to the degradation of released factors. Accordingly, all analyses in this study focused on days 1-3 post-infection. Previous work suggested that Mtb infection induces a cell death pathway dependent upon type I IFN signaling [49]. To test if this pathway is engaged in THP-1 cells, we tested the efficacy and duration of IFNAR neutralization by stimulating THP-1 cells with IFN-β in the presence or absence of IFNAR-neutralizing antibodies, followed by immunoblotting for phosphorylated STAT1 (pSTAT1) and total STAT1 (Fig. S1A–D). The results confirmed that STAT1 is activated during Mtb infection in THP-1 cells, and that IFNAR neutralization effectively inhibits STAT1 phosphorylation (Fig. 1C, D). Across four independent experiments, the ratio of pSTAT1 to total STAT1, as well as total STAT1 levels, did not significantly change (Fig. S1E, F). Despite effective IFNAR blockade, no significant reduction in LDH release was observed in Mtb-infected THP-1 cells (Fig. 1E), indicating that IFNAR inhibition does not attenuate cell death in them. Furthermore, ELISA analysis revealed that while IFN-β is produced during Mtb infection, the absolute levels are low. These findings collectively suggest that type I IFNs are not actively involved in THP-1 cell death during *ex vivo* Mtb infection (Fig. S1G).

**Fig. 1:**
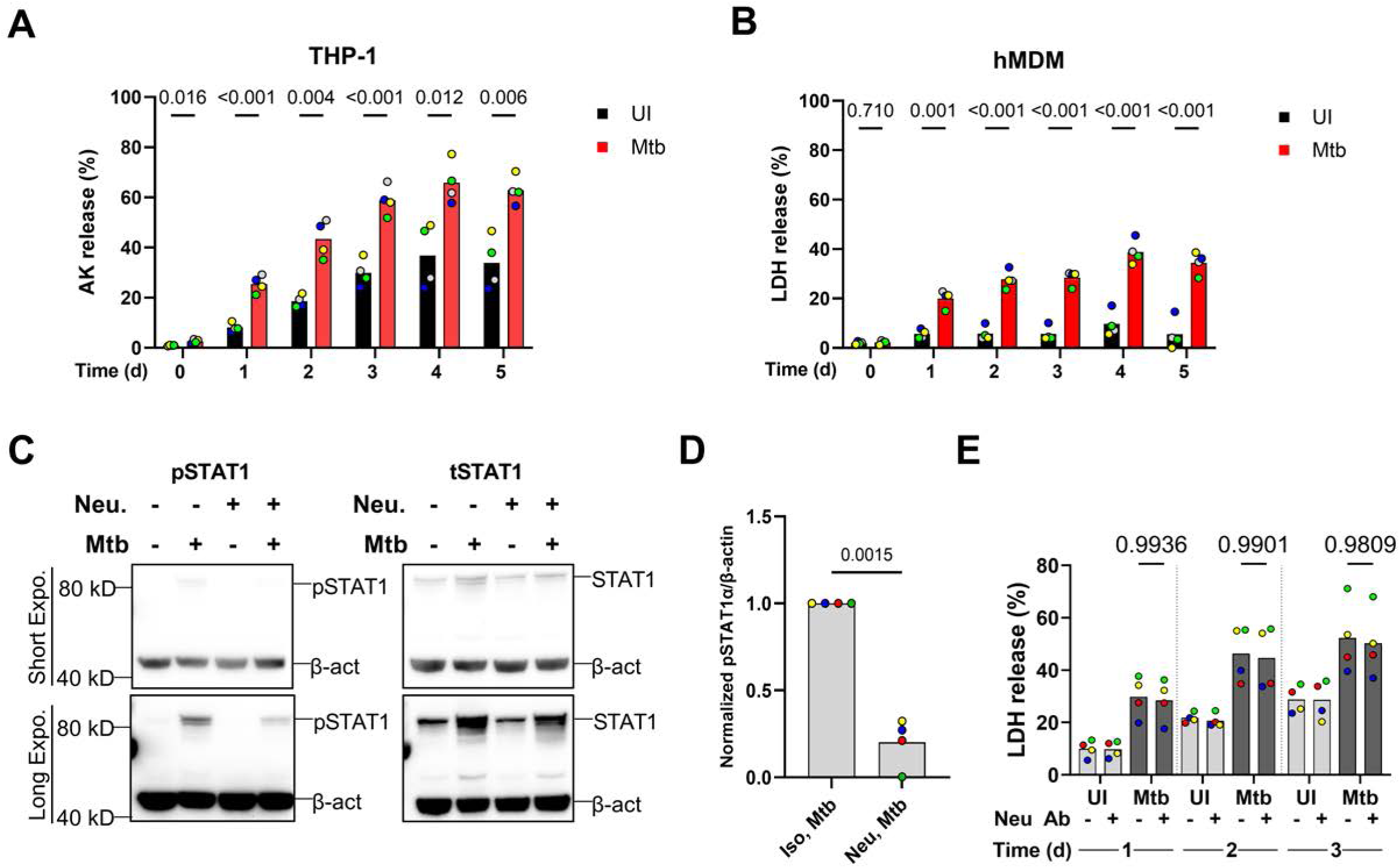
Kinetics of Mtb-mediated cell death induction and evaluation of IFN-β involvement. THP-1 cells (**A**, **C**, **D**, **E**) and hMDMs (**B**) were infected with H37Rv Mtb at an MOI of 5. **A** Mean percentages of AK release relative to the positive control. Each colored dot represents an independent experiment (N = 4). **B** Mean percentages of LDH release relative to the positive control. Each dot represents an independent experiment (N = 6), with colors indicating distinct human monocyte donors (N = 3). **C** Representative western blot images showing STAT1 expression and phosphorylation following Mtb infection and IFNAR neutralization. Data are representative of four independent experiments (N = 4). **D** Quantification of band intensities shown in panel C. p-STAT1α intensity was normalized to β-actin from the same lane. **E** Mean percentages of LDH release relative to the positive control. Each colored dot represents an independent experiment (N = 4). Welch’s t-tests were performed for panels **A**, **B**, and **D**; *p*-values are shown on the graphs. Two-way ANOVA was performed for panel **E**, with *p*-values presented accordingly.

### Mtb induces minor apoptosis in THP-1 cells, followed by secondary necrosis. Mtb does not induce apoptosis in hMDMs

To determine whether Mtb induces apoptosis in THP-1 cells or hMDMs, YO-PRO-1 (YO) staining and western blot (WB) analyses were performed. Early apoptotic cells are expected to be stained by YO but not Draq7, whereas necrotic cells are stained by both dyes. MG132-treated THP-1 cells exhibited an apoptotic phenotype and served as a positive control. These cells were stained and subjected to time-lapse live-cell imaging to validate that YO and Draq7 accurately discriminate early apoptotic and necrotic cells (Fig. S2A, also see supplementary video (SV) 1). THP-1 cells or hMDMs were then infected with dsRed H37Rv Mtb at an MOI of 5 and stained with YO and Draq7. Fluorescence microscopy showed that more than 90% of cells were infected at 1, 2, and 3 d post-infection (Fig. S3B and D), whereas fewer than 6% of total cells exhibited an early apoptotic phenotype (Fig. 2A and B; Fig. S3A and C).

**Fig. 2:**
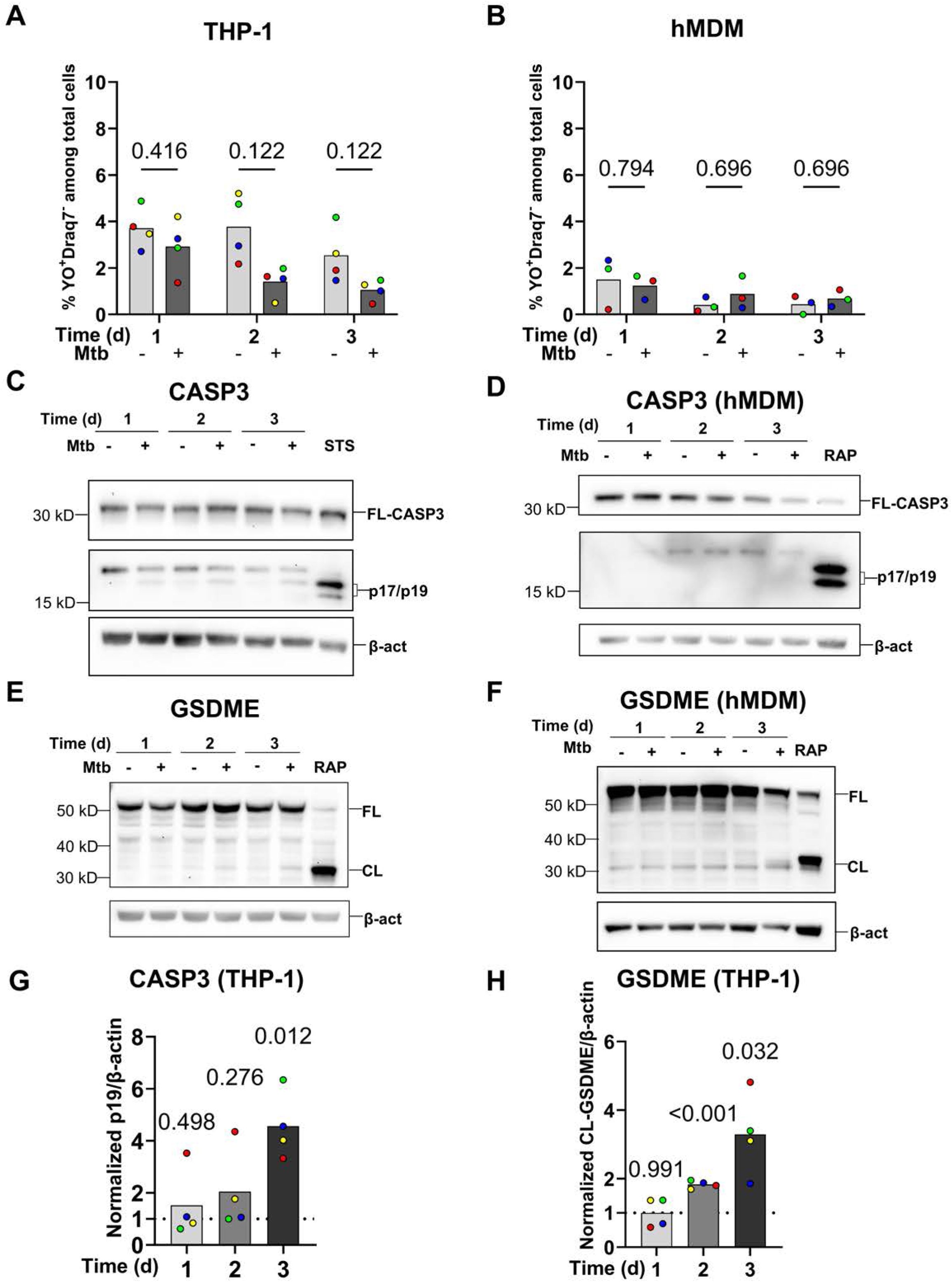
Mtb H37Rv does not induce apoptosis in THP-1 and hMDM but induces CASP3 and GSDME activation in THP-1 cells but not in hMDMs. **A, B** Percentages of early apoptotic cells among total cells assessed by fluorescence microscopy. THP-1 cells or hMDMs were infected with dsRed-expressing H37Rv Mtb at MOI:5 for 1–3 d, stained with YO-PRO-1 (YO) and Draq7, and imaged at 1, 2, or 3 d post-infection. More than 200 cells were analyzed per time point in each replicate (N = 3). Multiple paired t-tests were performed to compare infected groups with uninfected groups. and *p*-values are shown in the graphs. Each colored data point represents an independent experiment (**A**, THP-1, N = 4) or a distinct donor (**B**, hMDM, N = 3). **C–H** THP-1 cells were infected with w.t. H37Rv Mtb at MOI:5 for 1, 2, and 3 d or left uninfected. **C–F** Representative western blot images showing the cleavage of CASP3 (**C**, **D**) and GSDME (**E**, **F**) in THP-1 cells (**C**, **E**) or hMDMs (**D**, **F**), respectively. FL, full-length target protein; CL, cleaved target protein. STS, Staurosporine (1 μM, 6 h); RAP, Raptinal (10 μM, 6 h). **G, H** Bar graphs showing the quantification of normalized band intensities corresponding to THP-1 cleaved CASP3 (**G**), and GSDME (**H**). Each value in the infected groups was normalized to its uninfected counterpart from the same experiment. Band intensities were quantified using ImageLab software. One-sample t-tests were performed to compare the normalized values to 1, and *p*-values are shown in the graphs. Each colored data point represents an independent experiment (N = 4).

To further assess activation of apoptotic signaling pathways during Mtb infection, CASP-3 and GSDME were examined by WB. In THP-1 cells, cleaved CASP-3 and GSDME were readily detected, indicating significant activation (Fig. 2C, E, G, and H). In contrast, activated CASP-3 or GSDME bands were not detected in hMDMs (Fig. 2D and F). In THP-1 cells, activation of the upstream initiator caspases CASP-8 and CASP-9 was also analyzed by WB (Fig. S4A–D), revealing no detectable activation during Mtb infection. Collectively, these data indicate that, in Mtb-infected THP-1 cells, a subset of cells undergoes secondary necrosis, whereas early apoptotic events are infrequent and difficult to capture. In hMDMs, apoptotic cells are rare (∼2%) during Mtb infection and are detectable only by single-cell microscopic analysis.

### Analysis of pyroptosis and necroptosis after Mtb infection

Given that previous studies have demonstrated pyroptosis induction in various models during Mtb infection [29,33], we investigated whether pyroptosis constitutes a major form of Mtb-induced cell death in macrophages. THP-1 cells and hMDMs were infected with H37Rv Mtb at MOI:5 for 4 h, and cell lysates were collected after 1, 2, and 3 d post-infection. Western blot analysis revealed significant GSDMD cleavage in THP-1 cells following Mtb infection, whereas no detectable cleavage was observed in hMDMs (Fig. 3A–D). Notably, the blots show two bands in the 30–32 kDa region (Fig. 3A). Fig. S5A demonstrates that both bands are detected using two distinct anti-GSDMD antibodies. Subsequent infections with multiple mycobacterial strains showed that all tested strains induced GSDMD cleavage and cell lysis, resulting in the same two bands (Fig. S5B–E). The positive control confirmed that the upper band corresponds to the cytotoxic, cleaved form of GSDMD, which is the focus of our study. Quantitative analyses suggest that different mycobacterial strains may induce alternative forms of GSDMD cleavage (Fig. S5C, D). However, since the lower band was consistently present across all infected conditions, it is unlikely to be specific to Mtb virulence. The ELISA (Fig. S5F) results also demonstrate that THP-1 cells secrete significant amounts of IL-1β during Mtb infection. It is well-established that the secretion of IL-1β from Mtb-infected cells *ex vivo* is dependent on inflammasome components and activity [28,30,58], although one study showed that the secretion of IL-1β from Mtb-infected cells *ex vivo* is independent of GSDMD [59]. Both our previous findings [33] and the data presented in this study (Fig. 8J) indicate that IL-1β secretion from infected THP-1 cells is mediated by the non-canonical inflammasome CASP4/5. Therefore, we infer that the detected IL-1β secretion is associated with, and serves as an indicator of, pyroptosis induction. GSDMD can permeabilize mitochondria, triggering ROS release and subsequent necroptosis marked by MLKL phosphorylation [41]. Despite clear GSDMD activation in THP-1 cells, we did not observe MLKL phosphorylation at any time point. Similarly, in hMDMs, neither pyroptosis nor necroptosis was detected under the experimental conditions (Fig. 3H–K, S5K, L). The specificity of the pMLKL antibody was validated using HT-29 cells, which are highly sensitive to necroptosis induction (Fig. 3G–J). Necroptosis inducers robustly activated canonical necroptotic signaling and induced MLKL phosphorylation in HT-29 cells, whereas no changes in MLKL phosphorylation were detected in THP-1 cells or hMDMs (Fig. 3H–J).

**Fig. 3:**
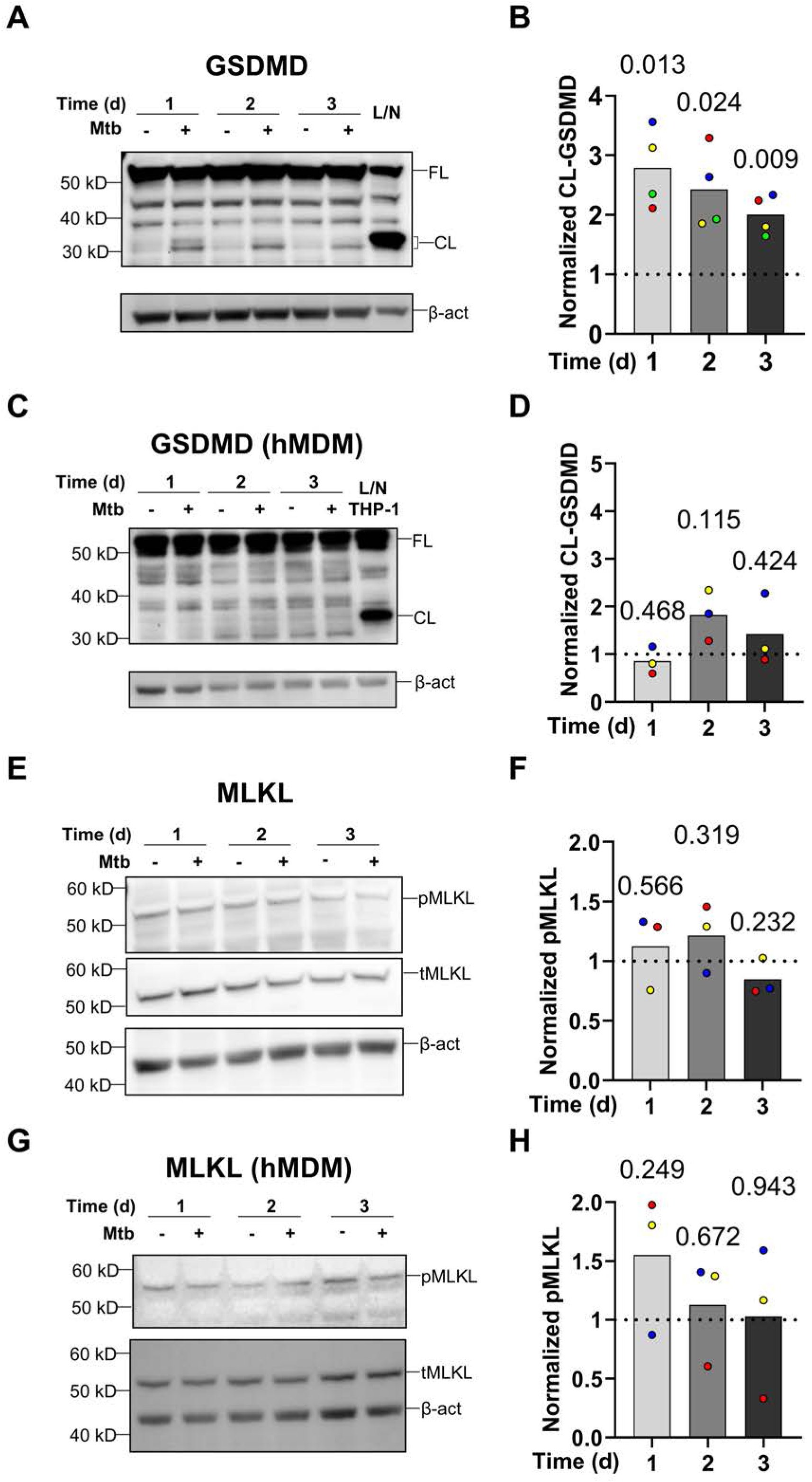
H37Rv Mtb induces GSDMD cleavage in THP-1 cells but not in hMDMs and does not induce MLKL phosphorylation in either cell type. THP-1 cells (**A**, **B**, **E**, and **F**) or hMDMs (**B**, **D**, **F**, and **H**) were infected with w.t. H37Rv Mtb at MOI:5 for 1-3 d or left uninfected. **A, C** Representative western blot images showing GSDMD cleavage in THP-1 cells (**A**) and hMDMs (**C**). **E, G** Representative western blot images showing MLKL phosphorylation in THP-1 cells (**E**) and hMDMs (**G**). FL, full-length target protein; CL, cleaved target protein. L/N, LPS (100 ng/mL, 4.5 h) plus nigericin (20 μM, 30 min) treatment, as a positive control. **B, D, F,** and **H** Quantification of band intensities corresponding to cleaved GSDMD (**B** and **D**) or phosphorylated MLKL (**F** and **H**), as indicated next to each Western blot. Band intensities were quantified using ImageLab software, normalized to the corresponding β-actin bands, and then expressed relative to the uninfected counterparts from the same experiment. Bar graphs display the normalized band intensities of the infected groups. One-sample t-tests were used to compare normalized values to 1, and *p*-values are shown in the graphs. Each colored data point represents an independent experiment (N = 4 or 3) for THP-1 cells (**B** and **F**) or an individual human donor (N = 3) for hMDMs (**D** and **H**).

### Mtb does not induce GPX4-regulated ferroptosis in THP-1 cells or hMDMs during the first three days after infection

Ferroptosis is a programmed cell death pathway that is activated after Mtb infection in BMDMs and during in vivo infections [42–44]. However, as most supporting studies have been conducted using murine *in vivo* or *ex vivo* models, we investigated whether ferroptosis occurs in human macrophage models. Immunoblotting analysis of GPX4 revealed that Mtb infection did not reduce GPX4 levels in either THP-1 cells or hMDMs at 1–3 days post-infection (Fig. 4A–D). Lipid peroxidation is a hallmark of ferroptosis [60]. We performed Liperfluo (LF) staining followed by fluorescence microscopy in Mtb-infected THP-1 cells to image lipid peroxidation. Although the cell membrane permeability stain, Draq7, indicated a trend toward increased cell death in Mtb-infected cells at 4- and 24-hour post-infection, LF fluorescence intensity remained unchanged, suggesting no significant involvement of lipid peroxidation in the cell death induction pathway (Fig. 4E–I). Ferrostatin-1 is a known ferroptosis inhibitor and was also tested in Mtb-infected THP-1 cells. LDH assay was performed to evaluate whether this ferroptosis inhibitor could reduce Mtb-induced cell death at early time points (Fig. 4J, K). However, ferrostatin-1 did not significantly reduce LDH release. These data collectively suggest that ferroptosis does not contribute to Mtb-induced cell death in human macrophages at the time points analyzed (Fig. 4E–I). Corresponding nested plots of LF fluorescence intensity are shown in Fig. S6A, B.

**Fig. 4:**
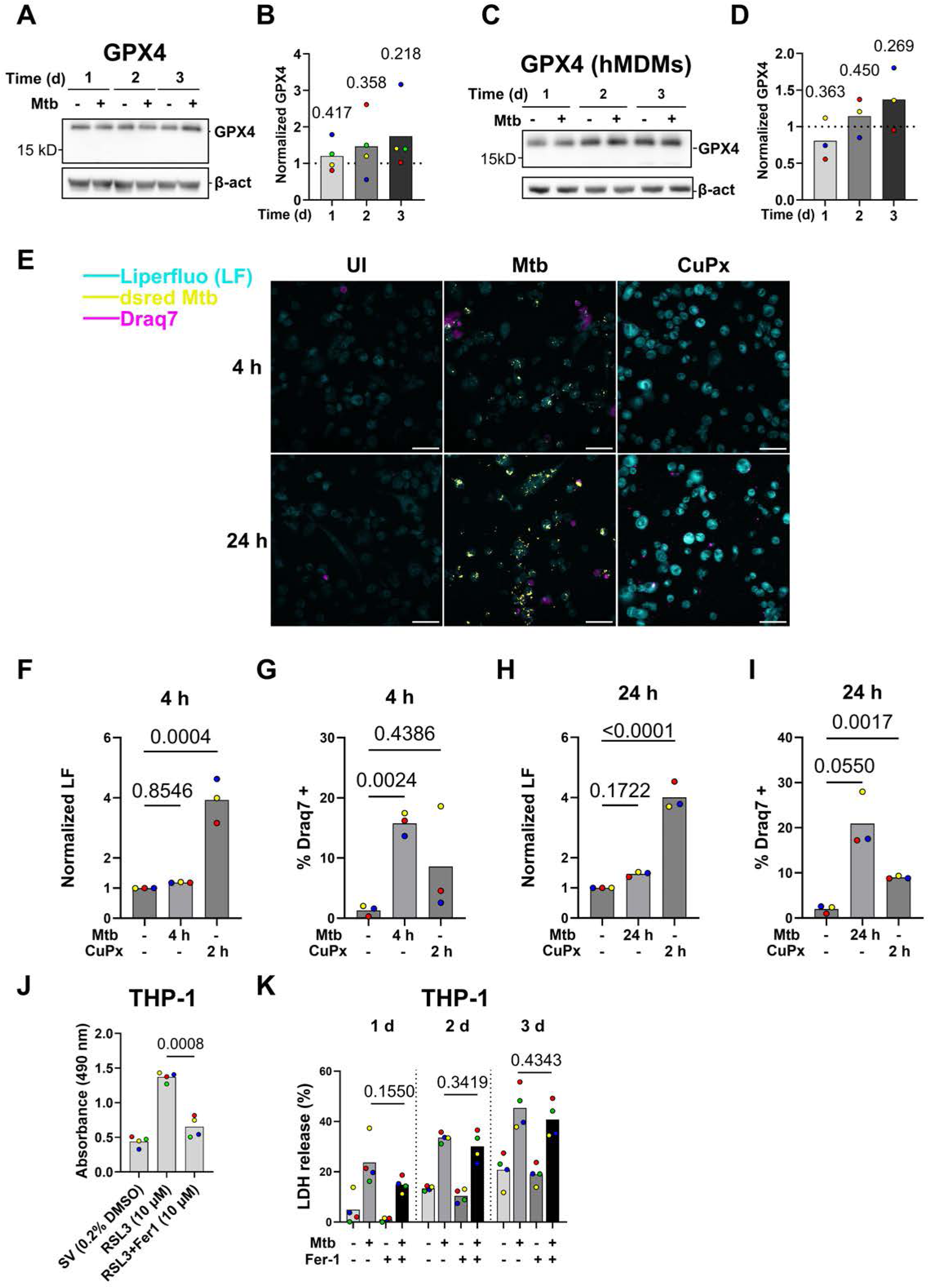
Mtb does not induce GPX4-regulated ferroptosis in THP-1 cells or hMDMs at the examined time points. **A, C** THP-1 cells or hMDMs were infected with H37Rv Mtb at MOI:5 for 1-3 d or left uninfected. Representative western blot images show the GPX4 levels in THP-1 cells (**A**) (N = 4) or hMDMs (**C**) (N = 6). **B, D** Band intensities of GPX4 were quantified using ImageLab software, normalized to the corresponding β-actin bands, and then expressed relative to the uninfected counterparts from the same experiment. Bar graphs display the normalized band intensities of the infected groups. One-sample t-tests were used to compare normalized values to 1, and *p*-values are shown in the graphs. One-sample t-tests were performed to compare normalized values to 1, and *p*-values are shown in the graphs. Each colored data point represents an independent experiment (N = 4) for THP-1 cells (**B**) or an individual human donor (N = 3) for hMDMs (**D**). **E** Representative confocal microscopic images showing the level of lipid peroxidation in THP-1 cells (N = 3). Cells were infected with dsRed H37Rv Mtb (yellow) at MOI:5 for 4 or 24 h, respectively, treated with cumene hydroperoxide (CuPx) for 2 h, or left uninfected and untreated. Staining was performed with Liperfluo (cyan) and Draq7 (magenta). Scale bars: 50 µm. **F–I** Quantification of normalized Liperfluo fluorescence intensities (**F** and **H**), or the percentage of Draq7-positive cells relative to the total cells in each field (**G** and **I**). Each dot represents the mean measurement from an independent experiment (N = 3), with 6 fields imaged per condition. Regions of interest (ROIs) were defined using Cellpose 2.0, and Raw Integrated Densities (RawIntDen) were calculated using FIJI. Nested One-way ANOVA (**F** and **H**) or One-way ANOVA with Brown-Forsythe correction (**G** and **I**) was performed; *p*-values are presented in decimal format. **J** LDH assay validating the efficacy of Ferrostatin-1. THP-1 cells were treated with solvent control (SV), RSL-3, or a combination of RSL-3 and Ferrostatin-1 (Fer-1) at the indicated concentrations. Background-corrected absorbance values for each group are shown. **K** LDH assay measuring necrosis in infected and uninfected THP-1 cells in the presence or absence of Fer-1. THP-1 cells were infected with w.t. H37Rv Mtb at MOI:5 for 1–3 d with or without Fer-1 (10 μM) (N = 4). The percentage of LDH release was calculated by normalizing background-corrected absorbance values to the lysis control.

### Mtb does not induce detectable lysosomal membrane permeabilization (LMP) in THP-1 cells for up to 2 hours prior to plasma membrane rupture

Lysosomal membrane permeabilization (LMP) is a well-characterized event that contributes to lysosome-dependent cell death (LDCD) [25]. Recent studies have reported LMP induction during Mtb-induced cell death in various experimental models [45,46,52]. In this study, time-lapse live-cell imaging was employed to monitor single Mtb-infected cells shortly before cell death. Lysoview 633 (LV) staining was used to assess LMP in THP-1 cells. To validate this approach, THP-1 cells were treated with L-leucyl-L-leucine methyl ester (LLoMe) and continuously imaged, with LV fluorescence intensity quantified (Fig. 5A, C; SVs 2, 3). A modified Chi-square analysis revealed a significant difference in LV signal compared to solvent-treated or untreated controls. Specifically, 81.25% of LLoMe-treated cells exhibited LMP, whereas only 10.25% of cells treated with DPBS did so (Fig. S7A–D), validating the experimental design and analytical approach. Following validation, the same procedure was applied to Mtb-infected THP-1 cells (MOI:2), which were imaged continuously for 20 hours. The modified Chi-square analysis indicated no significant difference in relative LV fluorescence intensity between infected cells and either bystander (Bys) or uninfected (UI) cells (Fig. 5B, D, SVs 4–6). Interestingly, bystander cells showed an increase in LV intensity, potentially reflecting lysosomal biogenesis. At the single-cell level, LMP was observed in 8.57% of infected cells, 8.45% of UI cells, and 2.44% of bystander cells (Fig. S7E–J). Collectively, these findings indicate that Mtb does not significantly induce LMP in THP-1 cells within 2 hours prior to PMR.

**Fig. 5:**
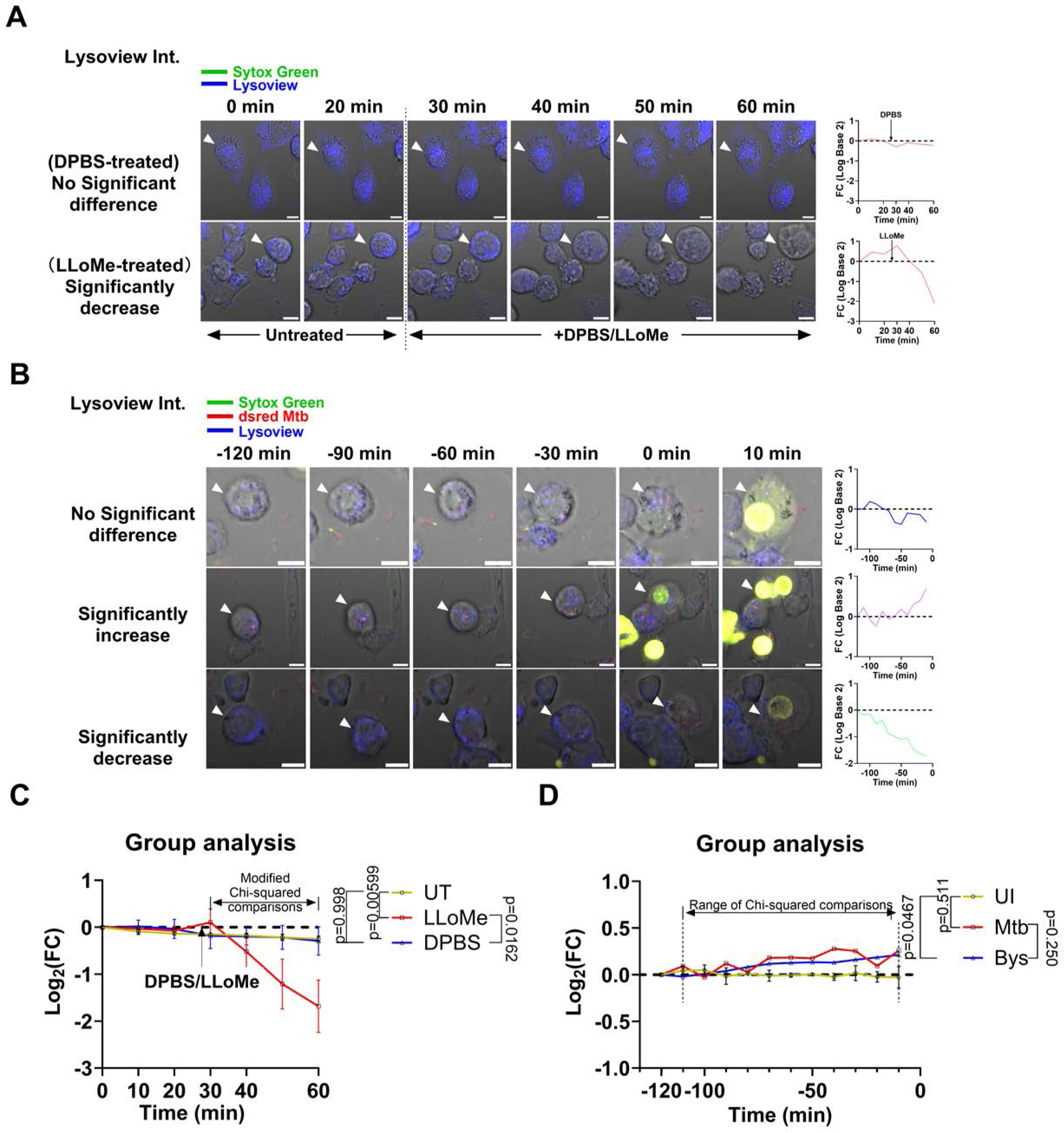
Mtb does not induce detectable lysosomal membrane permeabilization (LMP) in THP-1 cells 2 hours prior to cell death. **A** Representative time-lapse live-cell imaging of THP-1 cells showing changes in Lysoview (LV, blue) fluorescence intensity over time. Cells were stained with LV and Sytox Green (Syt G, green), then treated with LLoMe or DPBS approximately 25 minutes after imaging began. White arrowheads indicate tracked cells. **B** THP-1 cells were infected with dsred H37Rv Mtb at MOI:2 then stained with LV and Syt G. Representative time-lapse live-cell imaging of THP-1 cells show changes in LV fluorescence intensity over time. 0 min is defined as the time when a cell first becomes Syt G^+^. Negative time points indicate the duration (up to 120 min) before time point 0, during which LV intensities were quantified. The line graphs next to the confocal microscopic images display the logarithm (log) (base 2) of fold change (FC) over time (**A** and **B**). **C, D** corresponding to **A** or **B**, the line graphs show the log2 FC changing over time. In each independent experiment, 2 scenes were imaged for each condition, and 4 for the untreated (UT) (uninfected) condition. A total of 3 independent experiments were performed (N = 3). The UT dataset is the same as the UI from the infection experiments. 39, 48, and 71 cells were tracked and included in the statistical analysis for the DPBS, LLoMe-treated, and UT groups. For **D**, 4 biological replicates were conducted for the Mtb and bystander (Bys) conditions (N = 4), and 3 for the UI condition (N = 3). 35, 41, and 71 cells were tracked and analyzed for the Mtb, Bys, and UI groups. In **C** and **D**, each line represents mean; error bars indicate SD. Modified Chi-squared analyses were performed. *p*-values are shown in graphs.

### Mtb induces DNA release in THP-1 cells and hMDMs during infection

Time-lapse live-cell imaging performed for LMP analysis also revealed extensive DNA release following PMR (Fig. 6A, SVs 7, 8). The morphology of the observed DNA release resembles that of the previously described METosis [61,62]. To characterize the dynamics of DNA release, in both Mtb-infected and uninfected conditions, we quantified the proportion of DNA-releasing cells among all dying cells within the imaging frame. Approximately 60% of dead, Mtb-infected THP-1 cells exhibited DNA release, compared with ∼20% of dead uninfected cells (Fig. 6B). Notably, the overall percentage of necrotic cells relative to total cells did not differ significantly between infected and uninfected groups (Fig. 6C). On average, DNA release occurred approximately 106 minutes after cells became Sytox Green-positive, marking the onset of cell death (Fig. 6E). In hMDMs, time-lapse imaging demonstrated that 32% of dead cells released DNA and formed extracellular web-like structures (Fig. 6A, D, SVs 9, 10). During Mtb infection, approximately 22% of hMDMs underwent necrosis (Fig. 6F). Among ∼2600 analyzed uninfected hMDMs, only 10 necrotic cells were detected, of which 5 released DNA. Notably, in 3 of 6 independent repeats, no necrotic cells were observed. Zoomed-in images of these trap-like structures and the corresponding time-lapse videos are shown in Fig. S8A and SVs. 11 (THP-1) and 12 (hMDM). Suicidal NETosis induced by PMA in human neutrophils (hNeu) was imaged and used as one of the references to define criteria to distinguish DNA-releasing cells from cells without DNA release (Fig. S8B; SV 13). The criteria are described in the Materials and Methods section.

**Fig. 6:**
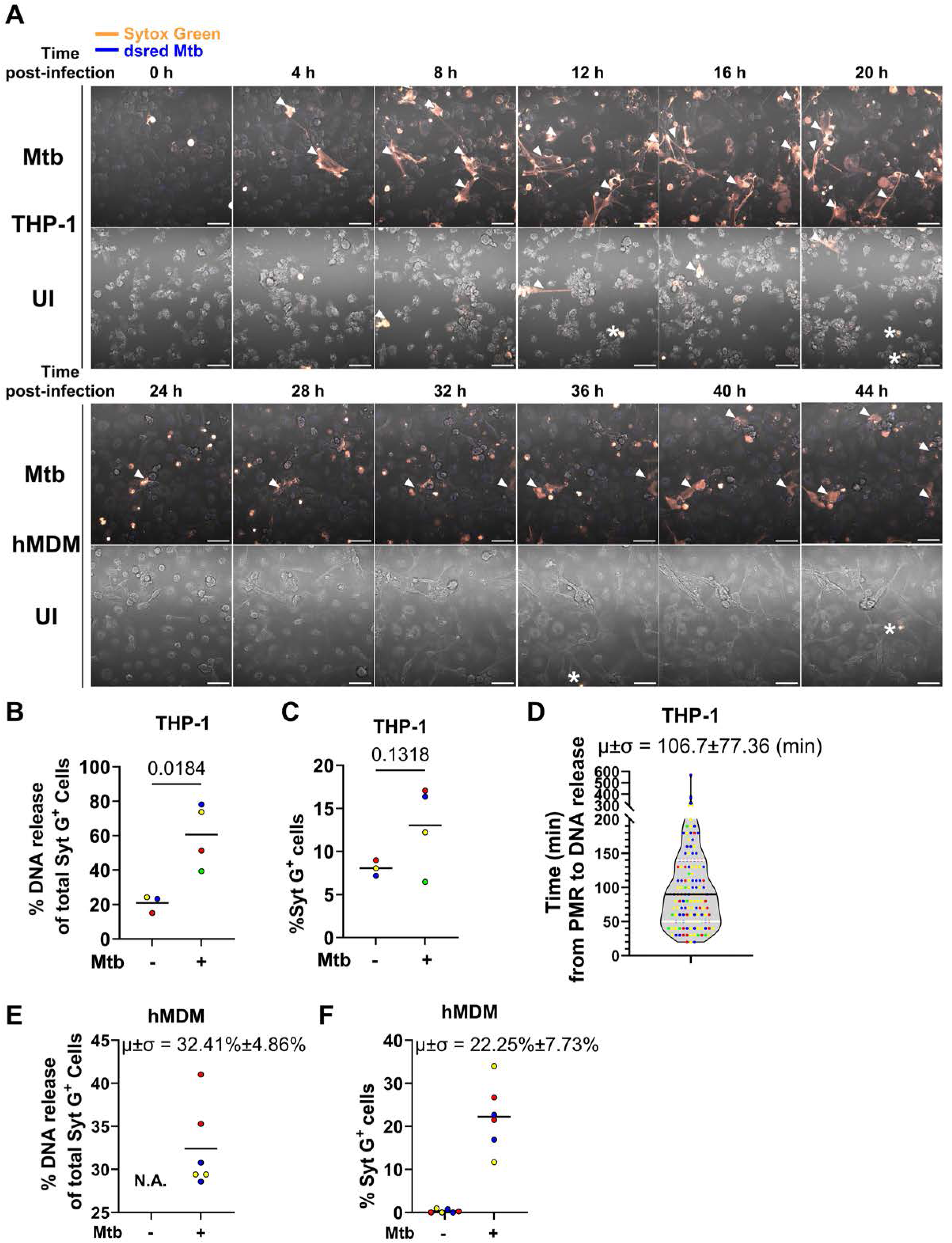
Mtb induces DNA release in THP-1 cells and hMDMs during the infection. **A**, THP-1 cell images extracted from the same time-lapse video shown in Fig. 5A and time-lapse confocal microscopic images of Mtb-infected hMDMs. THP-1 cells or hMDMs were infected with dsRed H37Rv Mtb (blue) at MOI:2 and then stained with Syt G (orange). The white arrowheads indicate extracellular, web-like DNA structures, while asterisks label cells that do not exhibit DNA release. Scale bars represent 50 μm. **B, C** Quantification of DNA release or plasma membrane rupture (PMR) ratios in uninfected and Mtb-infected THP-1 cells. DNA release events and total cell numbers in frame 1 of each video were quantified using FIJI software (the same for **E** and **F**). Each colored data point represents the average of 4 fields (N = 4 for Mtb; N = 3 for UI) and includes over 200 cells. Statistical analysis was performed using unpaired Welch’s t-tests, and *p*-values are shown in the graphs. **D** Quantification of the time duration from PMR to DNA release in Mtb-infected THP-1 cells. The number of frames between the two events was manually counted and converted to time (in minutes). The black dot line indicates the median, and the white dashed lines represent the 25th and 75th percentiles. Each dot represents a single trackable cell. The mean (μ) and standard deviation (σ) are indicated; the same notation is used hereinafter. Each color represents an independent experiment (N = 4). **E, F** Quantification of DNA release or PMR ratios in Mtb-infected or uninfected hMDMs. Each data point represents the average of 4 fields from an independent experiment (N = 6), with more than 200 cells analyzed per condition. Each color corresponds to a distinct human donor (N = 3). μ and σ are shown only for the infected groups. N.A., Not applicable.

### Markers of NETosis are detected on the extracellular DNA released by Mtb-infected THP-1 cells

Extracellular DNA released from Mtb-infected THP-1 cells and hMDMs exhibits a morphology resembling neutrophil extracellular traps (NETs) (Fig. 6A). NET formation and its associated cell death (NETosis), is a well-characterized form of PCD that conditionally involves histone citrullination and the activity of myeloperoxidase (MPO) and neutrophil elastase (NE) [63–65]. To investigate the mechanisms underlying DNA release from THP-1 cells during Mtb infection, a comparative analysis was performed between THP-1 cells and human neutrophils.

Immunofluorescence microscopy was used to assess the localization of MPO and citrullinated histone H3 (CitH3). Notably, the extracellular DNA released by Mtb-infected THP-1 cells was positively stained with antibodies against MPO and CitH3 (Fig. 7A–C, Fig. S9A), indicating their association with the DNA structures visualized by DAPI staining. However, compared with Mtb-infected neutrophils, the abundance of MPO and CitH3 on extracellular DNA from THP-1 cells was markedly lower (Fig. S9B, C). While neutrophils displayed considerable variability in MPO and CitH3 deposition on extracellular DNA, THP-1 cells consistently exhibited lower levels of these markers. Airyscan confocal microscopy analysis suggested that the extracellular DNA released from THP-1 cells could associate with extracellular Mtb bacilli (Fig. 7D, SV14). To assess the effect of extracellular DNA on THP-1 cells and Mtb, DNase I was used to treat uninfected THP-1 cells or THP-1 cells infected with w.t. H37Rv Mtb in the absence of gentamycin in the chase medium. DNase I in the culture medium effectively removed the trap-like structures formed by Mtb-infected THP-1 cells, as shown in Fig. S9E and F. DNase I treatment significantly reduced LDH release from Mtb-infected THP-1 cells by ∼10%, while it had no effect on the viability of uninfected THP-1 cells (Fig. 7E). CFU analysis indicated that DNase I treatment reduced extracellular bacilli by an average of ∼75% compared with untreated controls across 4 independent replicates (Fig. 7F). However, this reduction did not reach statistical significance. In addition, DNase I neither killed Mtb nor inhibited Mtb growth in 7H9 culture medium (Fig. S9D). Collectively, these findings suggest that extracellular DNA may modestly exacerbate cell death in infected or bystander cells during Mtb infection, while the trap-like structures do not directly kill extracellular Mtb.

**Fig 7:**
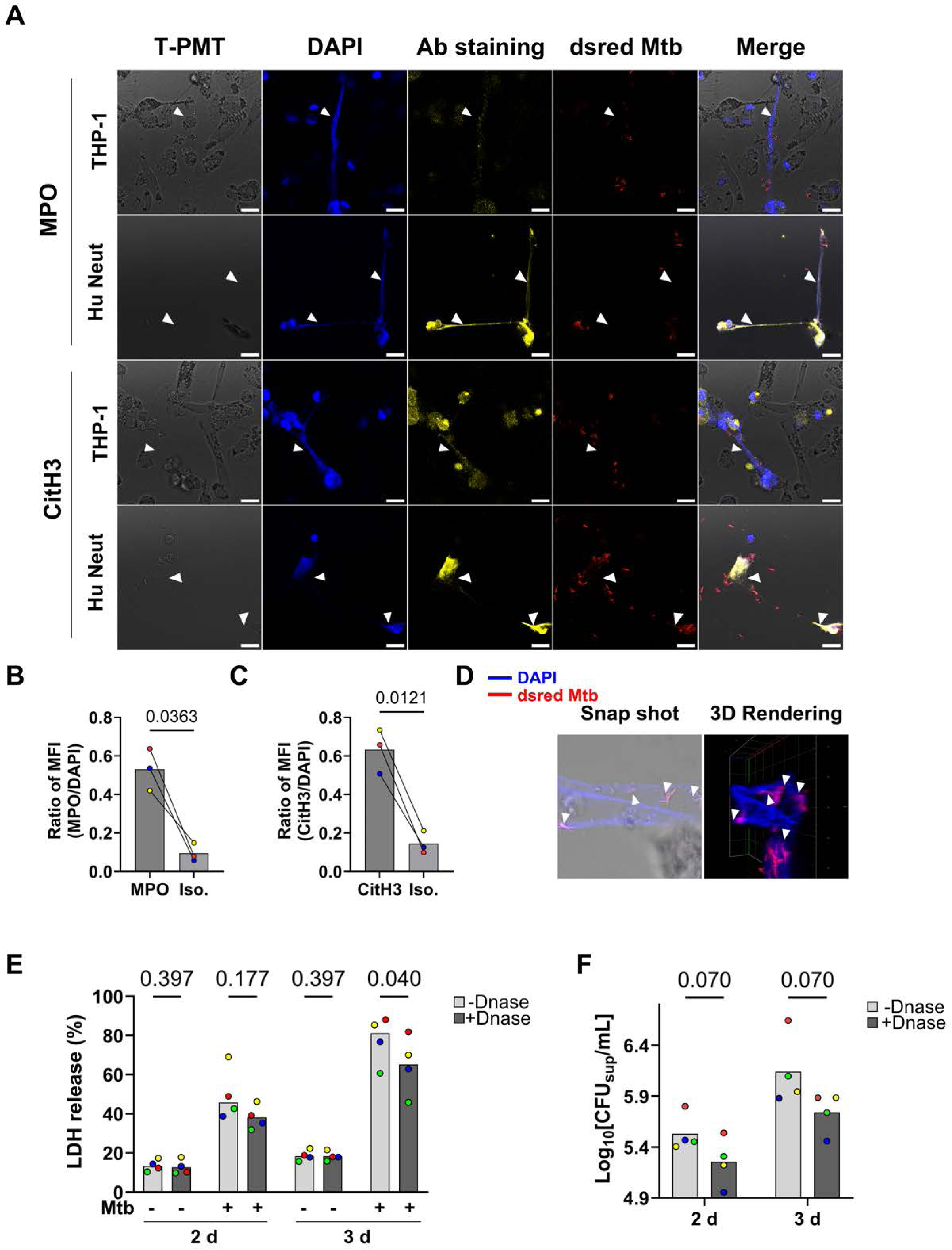
Markers of NETosis are detected on the extracellular DNA released by Mtb-infected THP-1 cells but that DNA has no bactericidal effect on extracellular bacteria. **A** THP-1 cells or human neutrophils (hNeu) (from 1 donor) were infected with dsred H37Rv Mtb (red) at MOI:2 for 20 hrs and fixed and stained with DAPI (blue) to visualize DNA. Alexa Fluor 647-conjugated antibodies (yellow) were applied to detect myeloperoxidase (MPO) or citrullinated Histone 3 (CitH3). White arrowheads indicate extracellular web-like DNA structures. Scale bars represent 50 μm. **B, C** Bar graphs showing the mean fluorescence intensity (MFI) ratios for DAPI and antibody staining on extracellular DNA. Intracellular fluorescence signals were manually excluded, and the MFI of DAPI or antibody staining on extracellular web-like DNA was quantified using FIJI. Paired t-tests were performed; *p*-values are reported in each graph. Each data point represents the average of 3-4 fields from an independent experiment, with each color denoting an independent experiment (N = 3). **D** Airyscan confocal image showing extracellular DNA released by infected THP-1 cells interacting with extracellular Mtb (N = 1). The left image is a 0.53 μm optical slice with merged signals from brightfield, DAPI (blue), and dsRed Mtb (red). 3D volume rendering (on the right) was performed using ZEISS Zen 3.6, showing the X-Y view. Each square in the background grid represents a 10 μm × 10 μm area. **E** LDH release assay showing the percentage of cell death in each group, calculated by normalizing absorbance values to the lysis control. THP-1 cells were infected with w.t. H37Rv Mtb at MOI:5 for 4 hours, washed with DPBS twice, and replenished with fresh medium. DNase I was added to half of the UI and infected groups at 100 U/mL, while the remaining groups were left untreated. Supernatants were collected at 2 and 3 d post-infection for LDH release analysis (**E**) and CFU enumeration (**F**) (N = 4 for both panels). **F** CFU analysis reflecting the number of viable extracellular Mtb bacilli. Supernatants from infected groups were harvested and plated on 7H11 agar plates. Samples were diluted 1:10 and 1:100 prior to plating, and only plates containing >20 colonies were used for quantification. Plates were monitored daily to ensure optimal colony size for counting, and CFUs were enumerated at 21 days post-plating.

### Pyroptosis signaling is not required for Mtb-induced cell death or DNA release

To identify potential regulators of extracellular DNA release, we investigated whether known cell death executioners are essential for this phenomenon.

Time-lapse live-cell imaging indicated that disulfiram, a GSDMD inhibitor, did not alter the extent of DNA release during Mtb infection (Fig. 8A, B, SVs 15, 16). Furthermore, disulfiram did not inhibit the induction of necrosis, as assessed by Syt G staining (Fig. 8C) and the LDH release (Fig. 8D). The control experiment shows that the disulfiram still effectively inhibited GSDMD activation in THP-1 cells after 20-hour treatment (Fig. 8E), and there is no difference in phagocytic capacity between the treated and untreated infected cells (Fig. S10A). These data indicate that GSDMD is not required for either Mtb-induced cell death or DNA release in THP-1 cells.

**Fig. 8:**
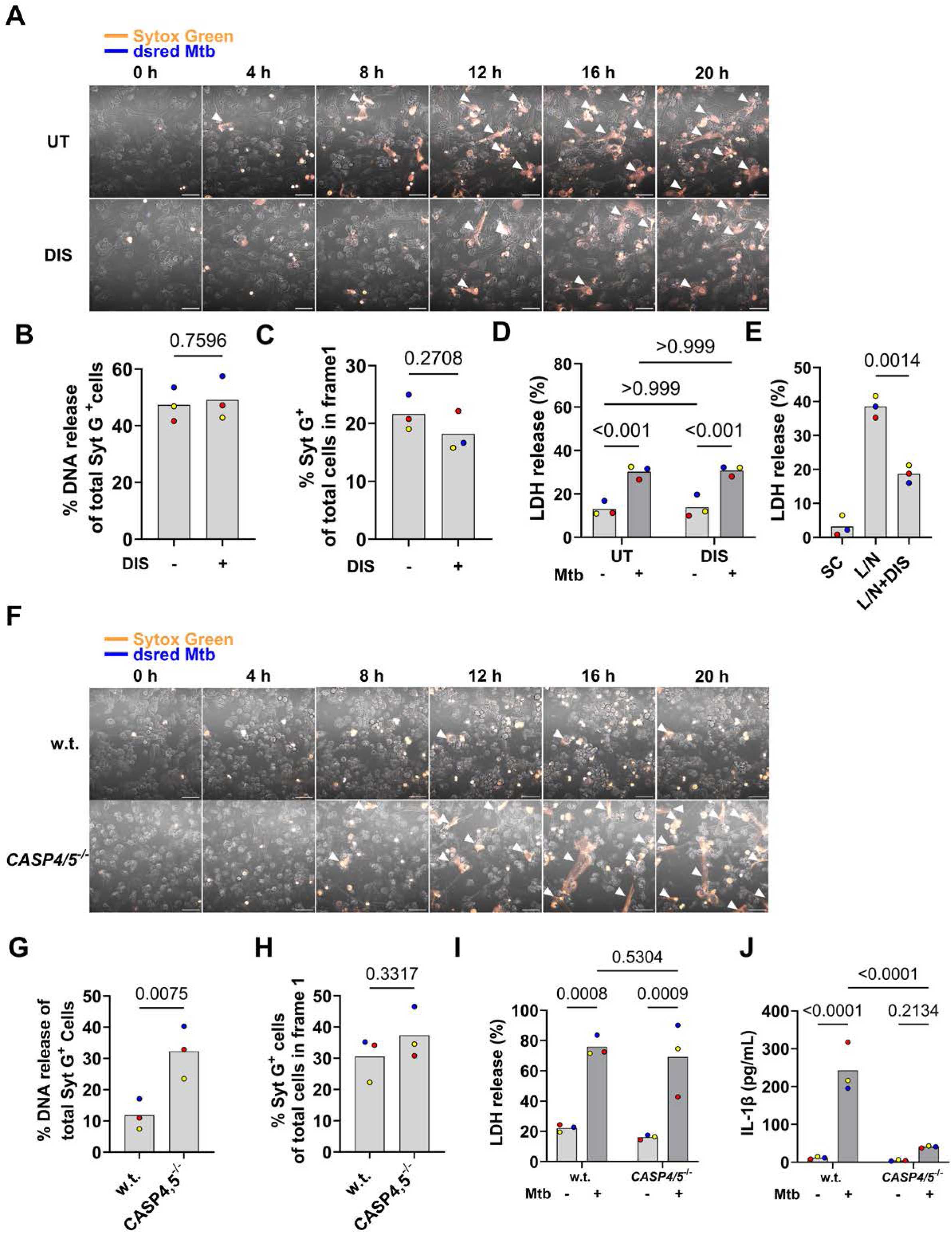
GSDMD and CASP4/5 are dispensable for Mtb-induced DNA release. **A** Confocal time-lapse images of THP-1 cells infected with dsRed H37Rv Mtb (blue, MOI: 2) and stained with Syt G (orange) for 20 h in the presence or absence of 20 μM disulfiram (DIS). UT, untreated. White arrowheads: extracellular DNA webs. **B, C** Quantification of DNA release events (**B**) and Syt G⁺ cells (**C**). **D, E** Percentages of LDH release relative to the lysis control. Validation of disulfiram efficacy (**E**): THP-1 cells were treated with 20 μM disulfiram prior to LPS and nigericin (L/N) stimulation, treated with L/N in the absence of DIS, or left untreated. SC, solvent control. **F** Confocal images of Mtb-infected *CASP4/5*^−/−^ and WT cells. THP-1 cells were a kind gift from Dr. Claire Bryant. **G, H** Quantification of DNA release events (**G**) and Syt G⁺ cells (**H**). **I, J** LDH, and IL-1β release profiles in *CASP4/5*^−/−^ cells. Data represent N = 3 independent experiments. *p*-values determined by Welch’s *t*-test (**B**, **C**, **E**, **F**, **H**) or two-way ANOVA (**D**, **I**, **J**). Scale bars: 50 μm.

Given the substantial pyroptosis observed in THP-1 cells upon Mtb infection, we next examined whether extracellular DNA release is a consequence of inflammasome activation. CASP4 and CASP5 have been recently reported to play critical roles in inflammasome activation in THP-1 cells during Mtb infection [33]. To assess their involvement, we employed *CASP4/5^−/−^* THP-1 cells. Time-lapse live-cell imaging demonstrated an increased frequency of DNA release events in *CASP 4,5^−/−^* cells compared to the wild-type THP-1 cells control (Fig. 8F and G; SVs 17, 18), despite comparable levels of cell necrosis between the two lines (Fig. 8H, I). WB confirmed gene deletion (Fig. S10B) and similar phagocytic capacity between the two cell lines (Fig. S10C), and ELISA revealed markedly reduced IL-1β secretion in *CASP 4,5^−/−^* cells during Mtb infection, consistent with previous findings (Fig. 8J) [45]. Collectively, these results indicate that CASP4/5-mediated inflammasome activation is associated with Mtb infection but is dispensable for the induction of necrosis and membrane rupture in THP-1 cells.

Previous studies have implicated NINJ1 in various forms of necrosis, pyroptosis, ferroptosis, and secondary necrosis [53,66–68]. Time-lapse live-cell imaging revealed no significant differences between *NINJ1^−/−^*and wild-type THP-1 cells in either the proportion of necrotic cells or the proportion of DNA release at 20 hours post-infection (Fig. S11A–D, SVs 19, 20). Genotype validation was performed by WB, and phagocytic capacity was assessed by fluorescence microscopy (Fig. S11F, G). Notably, *NINJ1^−/−^* THP-1 cells did not exhibit delayed membrane rupture during Mtb infection (Fig. S11D), but *NINJ1*^−/−^ THP-1 cells have a significantly higher level of LDH release during Mtb infection (Fig. S11E), suggesting that NINJ1 is involved in cell lysis but the initial PMR and DNA release in this context may occur through NINJ1-independent mechanisms.

## Discussion

In this study, we used THP-1 cells and hMDMs to investigate the modalities of cell death associated with Mtb infection. THP-1 cells and hMDMs undergo significant necrosis within the first 3 d post-infection (Fig. 1A, B). Previous work in murine macrophages shows that type I IFN signaling is involved in Mtb-mediated cell death induction [49]. Nevertheless, work by the same group demonstrated that Mtb infection of hMDMs and THP-1 cells did not induce substantial secretion of type I IFNs and that blocking type I IFN signaling did not affect cell death induction [48]. Consistent with this study, we show that IFN-β signaling is not involved in cell death signaling during the first three days of Mtb infection in THP-1 cells (Fig. 1E) and that Mtb infection results in a minimal increase in IFN-β production (Fig. S1G), which we also previously showed for hMDM [69]. These studies provide one example of the differences between murine and human macrophages regarding cell death pathways activated after Mtb infection.

Mtb limits the induction of host cell apoptosis to increase its virulence [8,16,17]. Consistently, the attenuated Mtb strain H37Ra induced an increased level of apoptosis compared to its virulent isogenic counterpart H37Rv [70,71]. Several Mtb proteins have been described to mediate the inhibition of host cell apoptosis [8,16]. For example, NuoG is a subunit of the type 1 NADH dehydrogenase of Mtb and a virulence factor involved in Mtb’s capacity to inhibit apoptotic cell death [19]. Mtb-infected THP-1 cells showed less than a 10% increase in apoptosis compared to uninfected controls, even after 5 days of infection [18,19,71]. Therefore, our findings of minimal (THP-1 cells) to no (hMDMs) induction of CASP3 activation (Fig. 2) align with these previous studies [18,19,71]. The absence of secondary necrosis induction, as demonstrated by little to no GSDME activation (Fig. 2G–J), further supports the lack of apoptosis induction during the timeframe of our study.

Our data reveal a divergence in inflammasome activation: while we observe the induction of GSDMD activation in THP-1 cells (Fig. 3A, B), we detect none in hMDMs (Fig. 3E, F). The IL-1β secretion from Mtb-infected THP-1 cells was detected (Fig. S2F). However, the concentration is substantially lower than that reported in other studies [30,59]. While differences in cell culture and differentiation conditions, as well as infection protocols, may contribute to this discrepancy, our findings suggest that pyroptosis is only modestly induced in THP-1 cells during Mtb infection. Phorbol 12-myristate 13-acetate (PMA), a standard agent for differentiating THP-1 cells, induces calcium influx. This increase in intracellular Ca²⁺ may potentiate inflammasome activation in THP-1 cells upon Mtb infection. Nevertheless, we let PMA-treated THP-1 cells rest for 2 days before infection, and so it is unlikely that the PMA treatment is the primary factor leading to increased inflammasome activation. Alternatively, THP-1 cells do not express the protein Sterile Alpha And TIR Motif Containing 1 (SARM1), a global inhibitor of inflammasomes present in human macrophages, which renders THP-1 cells more sensitive to inflammasome activation compared to hMDMs [72]. Interestingly, while there is some GSDMD activation in THP-1 cells (Fig.3 A, B), this does not seem to result in pyroptosis since there was no effect of caspase-4/-5 deletion on cell death (Fig. 8I). This result is consistent with our published data showing that the non-canonical inflammasome pathway is essential for inflammasome activation in THP-1 cells but does not affect cell death as measured by LDH [33].

Previous studies have reported extensive ferroptosis in Mtb-infected murine BMDMs (MOI:10, 1 d post-infection) and *in vivo* models [42,44]. Ferroptosis has also been observed in Mtb-infected hMDMs (MOI:10, 4 d post-infection) and human clinical samples [43,44]. Our data show that GPX4 levels are not markedly reduced during the first 3 days of Mtb infection and that no significant levels of lipid peroxidation could be detected (Fig. 4). This indicates that prolonged host–pathogen interaction or advanced TB disease progression may be required to detect significant ferroptosis in human macrophages. These findings suggest that ferroptosis may represent an early cell death modality in the context of Mtb infection in murine macrophages, but not in human macrophages.

Furthermore, several studies have reported LMP during Mtb infection [45,46,52]. However, our data show no evidence of LMP within the final 2 hours preceding plasma membrane rupture (Fig. 5). To the best of our knowledge, this is the first time that LMP was analyzed during Mtb infection at a single-cell level via live cell imaging. This type of analysis provides significantly higher resolution compared to bulk cell characterization via western blot or even snapshot analysis via fixed cell immunofluorescence microscopy. Differences between our results and the published data [45,46,52] could be due to: 1) the time of analysis, 2) host cell species, and/or 3) MOI. In other cellular contexts, LMP is frequently associated with apoptosis, pyroptosis, or ferroptosis [73–75], but our data suggest that LMP is not a major contributor to necrotic cell death in hMDMs during early Mtb infection.

One surprising finding of our work was the lack of engagement of any of the canonical cell death signaling pathways previously reported to be activated by Mtb. Our new data is consistent with data on Mtb-infected human macrophages which show that inhibitors of pyropotosis, necroptosis, apoptosis or ferroptosis do not inhibit necrotic cell death induction[76]. Our live cell imaging approach shows the release of DNA from dead Mtb-infected THP-1 cells (Fig. 6). Similar observations have been reported in Mtb-infected hMDMs, where METosis was claimed [74]. Notably, we are the first to demonstrate the dynamic process of DNA release from dying, Mtb-infected THP-1 cells and hMDMs using time-lapse live-cell imaging (Fig. 6A, SV 7, 9). This analysis revealed that Mtb induces plasma membrane rupture preceding DNA release, which contrasts with the vital NETosis observed during Mtb infection of neutrophils [77]. This approach also allowed us, for the first time, to quantify the relative importance of the DNA-release cell death phenotype compared to total cell death. We find that about 60% of all Mtb-infected THP-1 cells and about 30% of hMDMs deaths were associated with DNA release (Fig. 6B, C, E, and F). Additionally, we observed that the movement of surrounding, live macrophages appears to expand the extracellular DNA, forming a web-like structure (Fig. 6A, B, SV 7, 9), which could not be observed previously using microscopy approaches on fixed cells [61,62]. However, whether this DNA release in human macrophages represents a distinct cell death modality governed by a specific signaling pathway remains unclear. For instance, some studies have suggested that NETosis is closely linked to neutrophil GSDMD activation and pyroptosis [78]. Our findings indicate that all observed DNA release events occurred only after PMR and are independent of the proptosis signaling pathway (Fig. 8). Nevertheless, given the minimal presence of MPO and citH3 on the extracellular DNA (Fig. 7A–D), it is unlikely that the DNA release in THP-1 cells follows the NETosis signaling pathway. The DNase treatment assay demonstrates that eDNA does not kill Mtb but can exacerbate cell death in THP-1 cells, underscoring the biological relevance of our findings.

One of the limitations of the current study is that we have not determined if the DNA release observed is initiated via a novel and independent cell death signaling pathway or just a secondary event to a yet to be identified programmed necrosis pathway. The mechanistic understanding of such a pathway in human macrophages will be important for the development of new host-directed therapeutics. The consequence of extracellular DNA derived from macrophages and neutrophils is the potential engagement of TLR7 and TLR9 on bystander cells [79,80]. This activation can lead to an increase in type I IFN production, particularly by dendritic cells [80,81]. High type I IFNA production increases susceptibility of mice and humans to Mtb infections [82,83]. Consequently, blocking the signaling pathway leading to DNA release should be host protective.

## Acknowledgments

Purchase of the Zeiss LSM 980 Airyscan 2 was supported by Award Number 1S10OD025223-01A1 from the National Institute of Health. Amy Beaven (Imaging Core) for assistance with the use of the Zeiss LSM 980 Airyscan 2. We thank Zizheng Zhang, Xiangyu Liu (University of Maryland, College Park, MD, USA), and Junzheng Chu (Nankai University, Tianjin, China) for general guidance on Python programming. We acknowledge Dr. Katrin Mayer-Barber and Dr. Eduardo Amaral (NIAID) for their helpful discussions.

---

**Fig. S1: Optimization of the Interferon-alpha/beta receptor (IFNAR) neutralizing assay.**

**A** Immunoblot images showing STAT1 expression and phosphorylation in THP-1 cells following IFN-β treatment and IFNAR neutralization at various time points. Images are from a single experiment (N = 1). **B** Quantification of band intensities shown in **A**. **C** Western blot images showing STAT1 expression and phosphorylation in response to increasing concentrations of IFN-β and IFNAR neutralization. Images are from a single experiment (N = 1). **D** Quantification of band intensities shown in **C**. **E, F** Quantification corresponding to Fig. 1D, showing the relative ratio of p-STAT1α to STAT1 (**E**) and the normalized STAT1/β-actin ratio (**F**). For **F**, Grubb’s test identifies and excludes an outlier at α = 0.05. **G** THP-1 cells were infected with H37Rv Mtb at MOI: 5 for 1–3 days. IFN-β secretion was measured by enzyme-linked immunosorbent assay (ELISA). For **E**, **F**, and **G**, each colored dot represents an independent experiment (N = 4). Welch’s t-tests were used for statistical analysis, and *p*-values are shown on the graphs.

**Fig. S2: Validation of YO-PRO-1 and Draq7 staining in distinct cell death modalities.**

**A** Representative confocal fluorescence images of THP-1 cells treated with 10 μM MG132 (N = 1). Following MG132 addition, cells were stained with YO-PRO-1 (100 nM) and Draq7 (300 nM) for 15 min before initiating time-lapse live-cell imaging. Cells were maintained at 37 °C and 5% CO₂ and imaged continuously for 20 h. Z-stack images (11 slices at 1 μm intervals) were captured every 10 min. Regions containing cells exhibiting characteristic apoptotic or necrotic morphology were selected and enlarged. Fluorescence signals for YO-PRO-1 (cyan), Draq7 (magenta), and merged images with transmitted-light overlays are shown. Five representative time points are displayed for each condition. The necrotic-cell image layout is identical to that of the apoptotic-cell panel. Scale bars: 20 μm.

**Fig. S3: Assessing early apoptosis by YO-PRO-1 and Draq7 staining in Mtb-infected THP-1 cells.**

**A, C** Representative confocal fluorescence images of THP-1 (**A**) or hMDMs (**C**). THP-1 cells (N = 4) or hMDMs (N = 3) were infected with dsRed H37Rv Mtb at MOI:5 for 1, 2, or 3 d. Cells were stained with YO-PRO-1 and Draq 7 for 15 min at 37 °C prior to imaging. Z-stack images (11 slices at 1 μm intervals) were captured for each field of view (FoV). White triangles indicate necrotic cells, while asterisks indicate early apoptotic cells. Fluorescence signals for YO-PRO-1 (cyan), Draq7 (magenta), dsred (yellow), and transmitted-light signals are merged. Scale bars: 50 μm. **B, D** Quantification of percentages of infected cells among total cells for THP-1 cells (**B**) or hMDMs (**D**).

**Fig. S4: Mtb H37Rv does not induce CASP8 or 9 activation in THP-1 cells.**

**A, C** Representative western blot images showing the cleavage of CASP9 and CASP8, respectively. THP-1 cells were infected with H37Rv Mtb (+) at MOI:5 for 1-3 days or left uninfected (-). STS: Staurosporine (1 μM, 6 h). **B, D** Bar graphs showing the quantification of normalized band intensities corresponding to THP-1 cleaved CASP9 (**B**), and CASP8 (**D**). Each value in the infected groups was normalized to its uninfected counterpart from the same experiment. Band intensities were quantified using ImageLab software. One-sample t-tests were performed to compare the normalized values to 1, and *p*-values are shown in the graphs. Each colored data point represents an independent experiment (N = 3).

**Fig. S5: Mycobacterial species induce variable GSDMD cleavage, and different cell types have distinct sensitivity to necroptosis induction.**

**A** Western blot images showing GSDMD cleavage in THP-1 cells infected with w.t. Mtb H37Rv at MOI:5 for 1 d or treated with LPS and Nigericin (L/N) or left uninfected (UI). Blots were probed using CST and NOV antibodies. Images are from one experiment (N = 1). **B** THP-1 cells were uninfected or infected with *Mycobacterium smegmatis* (Msm), *Mycobacterium kansasii* (Mkan), and Mtb, all at MOI:5 or treated with L/N. Representative western blot images showing GSDMD cleavage induced by different mycobacterial strains in THP-1 cells (N = 4). **C, D** Quantification of normalized GSDMD cleavage band intensities corresponding to **B**. **E,** quantification of LDH release as a percentage of the lysis control, corresponding to **B**. Each colored data point represents an independent experiment (N = 4) for **C–E**. One-way ANOVA with Brown-Forsythe correction was performed, and *p*-values are shown in the graphs. **F** ELISA measurement of IL-1β secretion from Mtb-infected THP-1 cells (MOI:5) at 1, 2, and 3 d. Multiple unpaired t-tests with Welch’s correction were performed (N = 4). **G, H** Immunoblot analysis of MLKL phosphorylation during necroptosis induction or inhibition in different cell types. SV, solvent control (0.3% DMSO); TSZ, TNF-α (100 ng/mL, O/N), SM-164 (10 μM, O/N), and zVAD-FMK (20 μM, O/N); TSZN, TSZ plus necrostatin-1 (10 μM, O/N). **G,** HT-29 cells were treated with SV, TSZ, or TSZN (N = 1). Band intensities were quantified using ImageLab software and normalized to SV for each target protein; normalized values are shown above the corresponding bands. **H** THP-1 cells, hMDMs, or HT-29 cells were treated with SV or TSZ. Quantification of normalized band intensities corresponding to phosphorylated MLKL is shown. Values from TSZ-treated samples were normalized to their SV-treated counterparts from the same experiment. Band intensities were quantified using ImageLab software. One-sample t-tests were performed to compare normalized values to 1, and *p*-values are shown in the graphs. Each colored data point represents an independent experiment using THP-1 cells or a distinct human donor for hMDMs (N = 3). **K** and **L** Quantification of normalized band intensities corresponding to phosphorylated MLKL from Fig. 3F or **H**, respectively. One-sample t-tests were performed to compare normalized values to 1, and *p*-values are shown in the graphs. Each colored data point represents an independent experiment (N = 3) for THP-1 cells (Fig. 3F) or distinct human donors (N = 3) for hMDMs (Fig. 3H).

**Fig. S6. Nested plot showing normalized Liperfluo fluorescence intensities in THP-1 cells.**

**A, B** Corresponding to Fig. 4F and H, Nested plots displaying Liperfluo RawIntDen quantification in individual THP-1 cells. Each dot represents a single cell, and each column corresponds to an independent experiment (N = 3). The number of cells per experiment ranges from 339 to 656. Black bars indicate the mean fluorescence intensity for each experiment. Statistical comparisons were performed using nested one-way ANOVA, and *p*-values are presented in the graphs.

**Fig. S7: Mtb does not induce detectable lysosomal membrane permeabilization (LMP) in THP-1 cells 2 hours before cell death induction.**

**A-J** The log (base 2) of normalized Lysoview (LV) fluorescence intensity was plotted against time. The grey curves represent the mean of the UI/UT condition, which served as a baseline for comparison. Each data point indicates the fluorescence value at a specific time point for a single cell with each color representing an independent experiment (N = 3). Each curve represents the time course of a single cell. The black arrow marks the time of DPBS or LLoMe addition. The number of total individual cells tracked in each experiment is shown in the title of each graph. **B, D, H, I,** and **J** are subsets of the datasets shown in **A**, **C**, **E**, **F**, and **G**, respectively, highlighting individual curves that exhibit significant differences relative to the baseline. The proportion of descending curves relative to the total number in each category was calculated and reported as the percentage of LMP.

**Fig. S8: Zoomed-in images illustrating the structures of released or expelled DNA.**

**A** Zoomed-in images corresponding to Fig. 6A showing the structures of released DNA from Mtb-infected THP-1 cells or hMDMs. **B** Time-lapse live-cell imaging of PMA-treated human neutrophils (hNeu) (N = 1). Image acquisition began immediately after the addition of 50 ng/mL PMA to hNeu. White arrowheads indicate extracellular, web-like DNA structures, while asterisks label cells that do not exhibit DNA release. Scale bars represent 50 μm.

**Fig. S9: Isotype controls for anti-MPO and anti-CitH3 antibody staining and validation of DNase I efficacy.**

**A** Representative confocal immunofluorescence images of dsRed H37Rv Mtb-infected THP-1 cells and hNeu undergoing DNA release. Cells were infected with dsRed H37Rv Mtb (red) at MOI: 2 for 20 hours, then fixed and stained with DAPI (blue) to visualize DNA. Alexa Fluor 647-conjugated mouse or rabbit IgG isotype control antibodies (yellow) were used to assess background staining. White arrowheads indicate extracellular web-like DNA structures. Scale bars: 20 μm. **B, C** Quantification of the normalized fluorescence intensity (FI) ratios. Each dot represents a single extracellular DNA domain. A total of 9 or 10 domains from 3 independent experiments (N = 3) were included in the analysis. Statistical comparisons were performed using unpaired t-tests with Welch’s correction. One outlier from the hNeu group in **C** was identified and removed by the ROUT test (Q = 1%). **D** Comparisons of ODs between DNase I-treated and untreated Mtb 7H9 cultures. Well-grown Mtb cultures were diluted to OD_600_ = 0.2 from different initial ODs. DNase I was added to the cultures at 100 U/mL, and the cultures were shaken at 140 rpm at 37 °C. OD_600_ values were measured at 3 and 5 d post-DNase treatment (N = 4). For each repeat, values from DNase-treated cultures were normalized to their untreated counterparts. **E** Analysis of DNase I activity in THP-1 culture medium at different time points by agarose gel electrophoresis. DNase I was diluted in RPMI 1640 medium containing 10% FBS to 100 U/mL and incubated at 37 °C for 0, 1, 2, or 3 days. After the incubation, 10 μL of the pre-incubated DNase I solutions from all groups were mixed with 10 μg (1 μL) of shredded salmon sperm DNA and incubated at 37 °C for 1 hour. Then the reaction mixtures were mixed with loading buffer and analyzed by agarose gel electrophoresis (N = 1). **F** Confocal fluorescence images of Mtb-infected THP-1 cells pre- and post-DNase I treatment. THP-1 cells were infected with dsRed H37Rv Mtb (blue) at MOI 5. 4 h after bacterial addition (defined as time 0), cells were washed twice with DPBS and replenished with fresh culture medium. At 24 h post-infection, cells were stained with Syt G (orange) and imaged, with three random fields of view (FoVs) acquired. DNase I (100 U/mL) was then added to the same well, and cells were incubated for 1 h at 37 °C under 5% CO₂. Three additional FoVs from the same well were imaged post-DNase treatment using identical microscope settings (N = 1).

**Fig. S10: Comparison of phagocytic capacity among different cell groups and validation of *CASP4*/*5* genotype by immunoblotting.**

**A, C** Quantification of the percentage of Mtb internalization relative to the total cells in corresponding frames (N = 3). Welch’s t-tests were performed, and *p*-values are shown in the graph. **B** Immunoblot and Ponceau red staining confirming the genotype of THP-1 cell lines (N = 1). THP-1 cells were differentiated and left untreated.

**Fig. S11: NINJ1 promotes cell lysis of Mtb-infected THP-1 cells but is not required for DNA release.**

**A** Representative confocal time-lapse images showing Mtb-infected *NINJ1*^−/−^and w.t. THP-1 cells undergoing DNA release. *NINJ1*^−/−^ and its w.t. counterpart are kind gifts from Dr. Liron David. Cells were infected with dsRed H37Rv Mtb (blue) at MOI:2 for 20 hrs and stained with Syt G (orange) to visualize DNA. White arrowheads indicate extracellular web-like DNA structures. Scale bars represent 50 μm. **B, C** Quantification of the percentage of DNA release (**B**) and Syt G^+^ cells (**C**). Welch’s t-tests were applied for statistical analysis. **D** Quantification of time durations from Mtb internalization to PMR. A nested t-test was performed. **E** LDH release relative to the lysis control. Two-way ANOVA was applied for statistical analysis. **F** Immunoblot confirming *NINJ1* genotype in differentiated, untreated THP-1 cells (*N* = 1). LD, initials of Liron David. **G** Quantification of the percentage of Mtb internalization relative to the total cells in corresponding frames. Welch’s t-tests were performed. Experiments were repeated 3 times (N = 3), and *p*-values were reported in the graphs for **B**, **C**, **D**, **E**, and **G**.

