## Supplementary material for "Characterization of programmed cell death pathways activated in *Mycobacterium tuberculosis*-infected human macrophages": SupFigures.pdf

Fig. S1

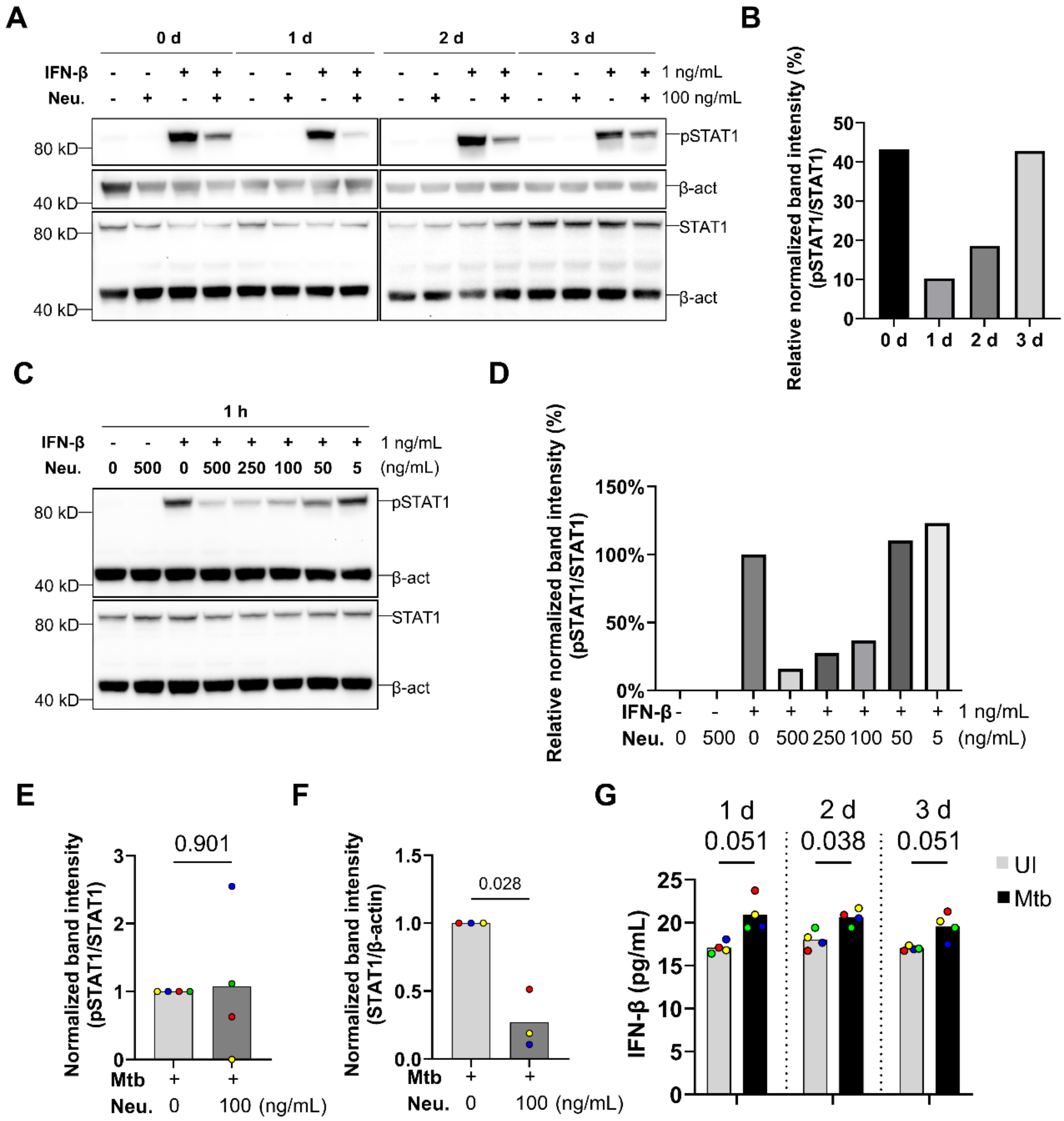
A
0 d
1 d
2 d
3 d
IFN-β
Neu.
1 ng/mL
100 ng/mL
80 kD
40 kD
pSTAT1
β-act
STAT1
B
Relative normalized band intensity (%) (pSTAT1/STAT1)
0 d 1 d 2 d 3 d
C
1 h
IFN-β
Neu. 0 500 0 500 250 100 50 5
1 ng/mL
(ng/mL)
D
Relative normalized band intensity (%) (pSTAT1/STAT1)
150%
100%
50%
0%
E
Normalized band intensity (pSTAT1/STAT1)
0.901
Mtb
Neu. 0 100 (ng/mL)
F
Normalized band intensity (STAT1/β-actin)
0.028
Mtb
Neu. 0 100 (ng/mL)
G
1 d 0.051
2 d 0.038
3 d 0.051
IFN-β (pg/mL)
UI
Mtb

Fig. S2

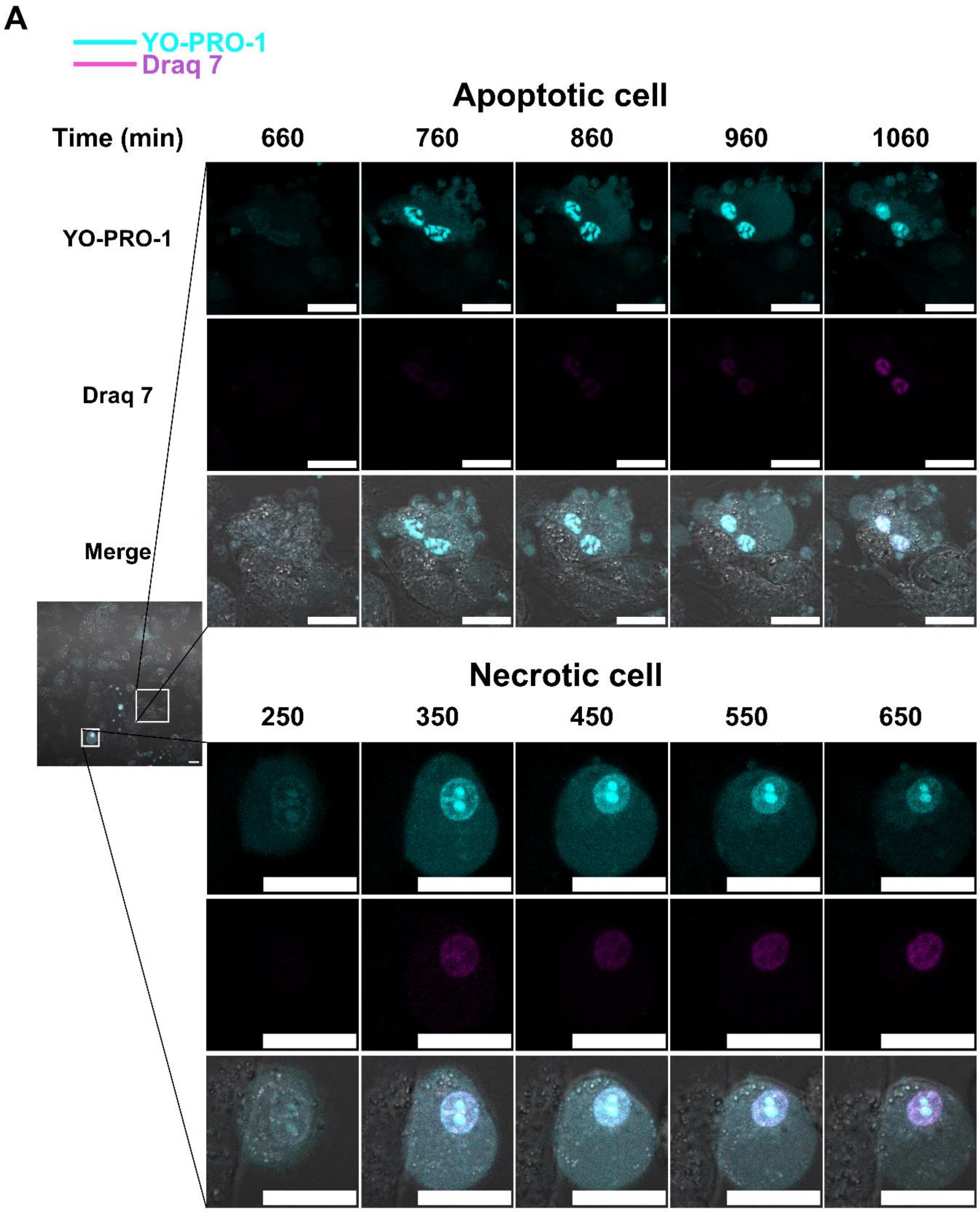
A
YO-PRO-1
Draq 7
Apoptotic cell
Time (min)
660
760
860
960
1060
YO-PRO-1
Draq 7
Merge
Necrotic cell
250
350
450
550
650

Fig. S3

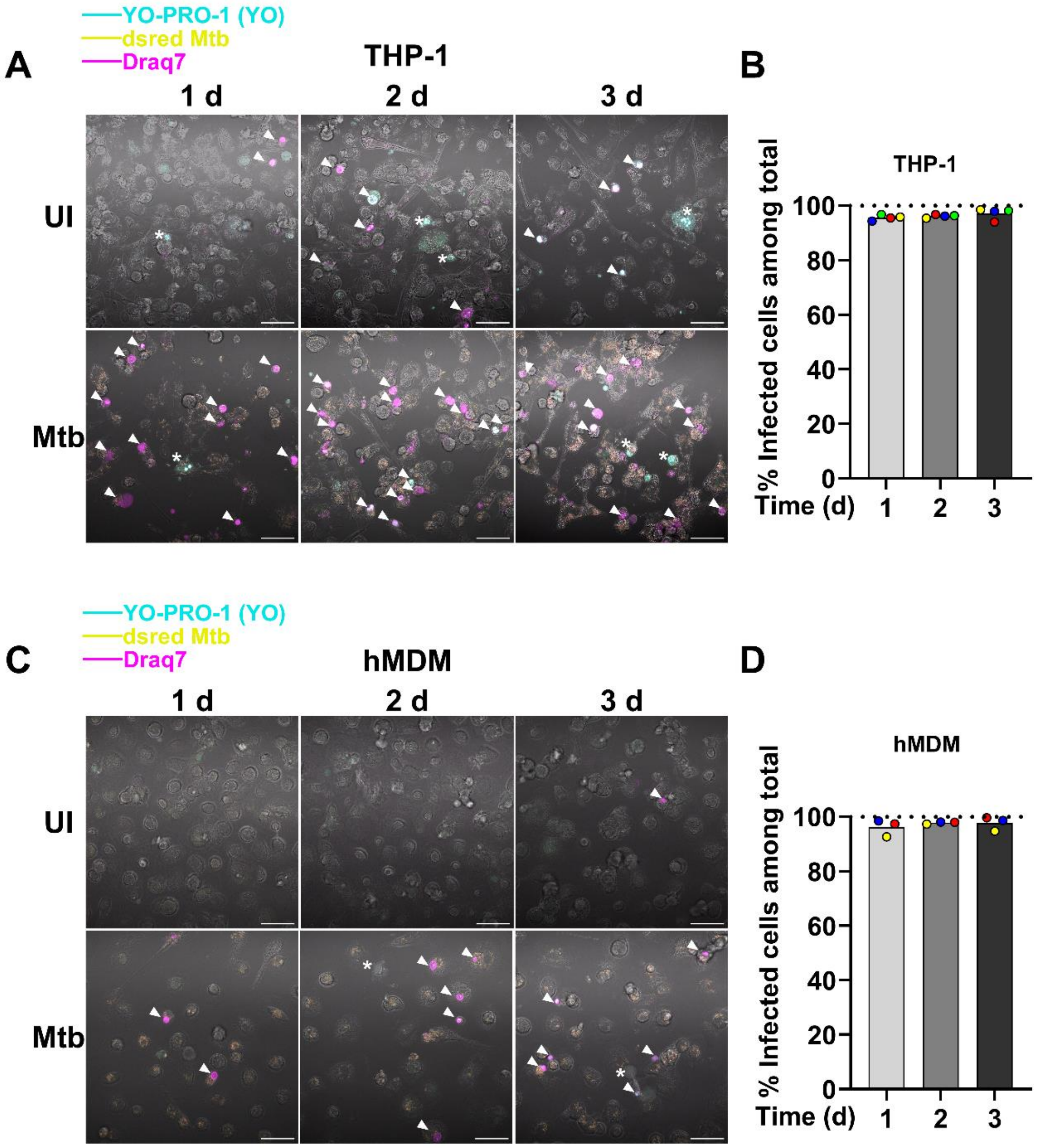

YO-PRO-1 (YO)
dsred Mtb
Draq7
A
THP-1
1 d
2 d
3 d
UI
Mtb
B
THP-1
% Infected cells among total
100
80
60
40
20
0
Time (d)
1
2
3
C
hMDM
1 d
2 d
3 d
UI
Mtb
D
hMDM
% Infected cells among total
100
80
60
40
20
0
Time (d)
1
2
3

Fig. S4

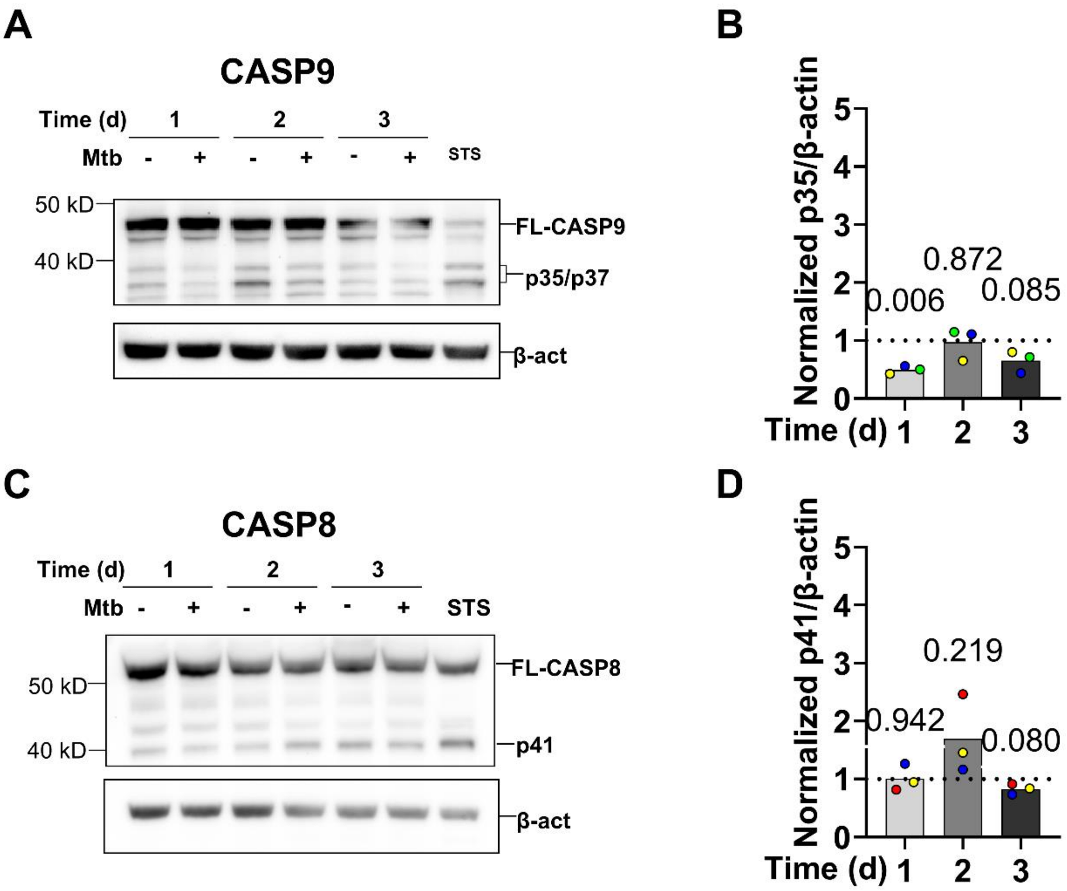

A
CASP9
Time (d) 1 2 3
Mtb - + - + - + STS
50 kD
40 kD
FL-CASP9
p35/p37
β-act
B
Normalized p35/β-actin
0.006
0.872
0.085
Time (d) 1 2 3
C
CASP8
Time (d) 1 2 3
Mtb - + - + - + STS
50 kD
40 kD
FL-CASP8
p41
β-act
D
Normalized p41/β-actin
0.942
0.219
0.080
Time (d) 1 2 3

### Fig. S5

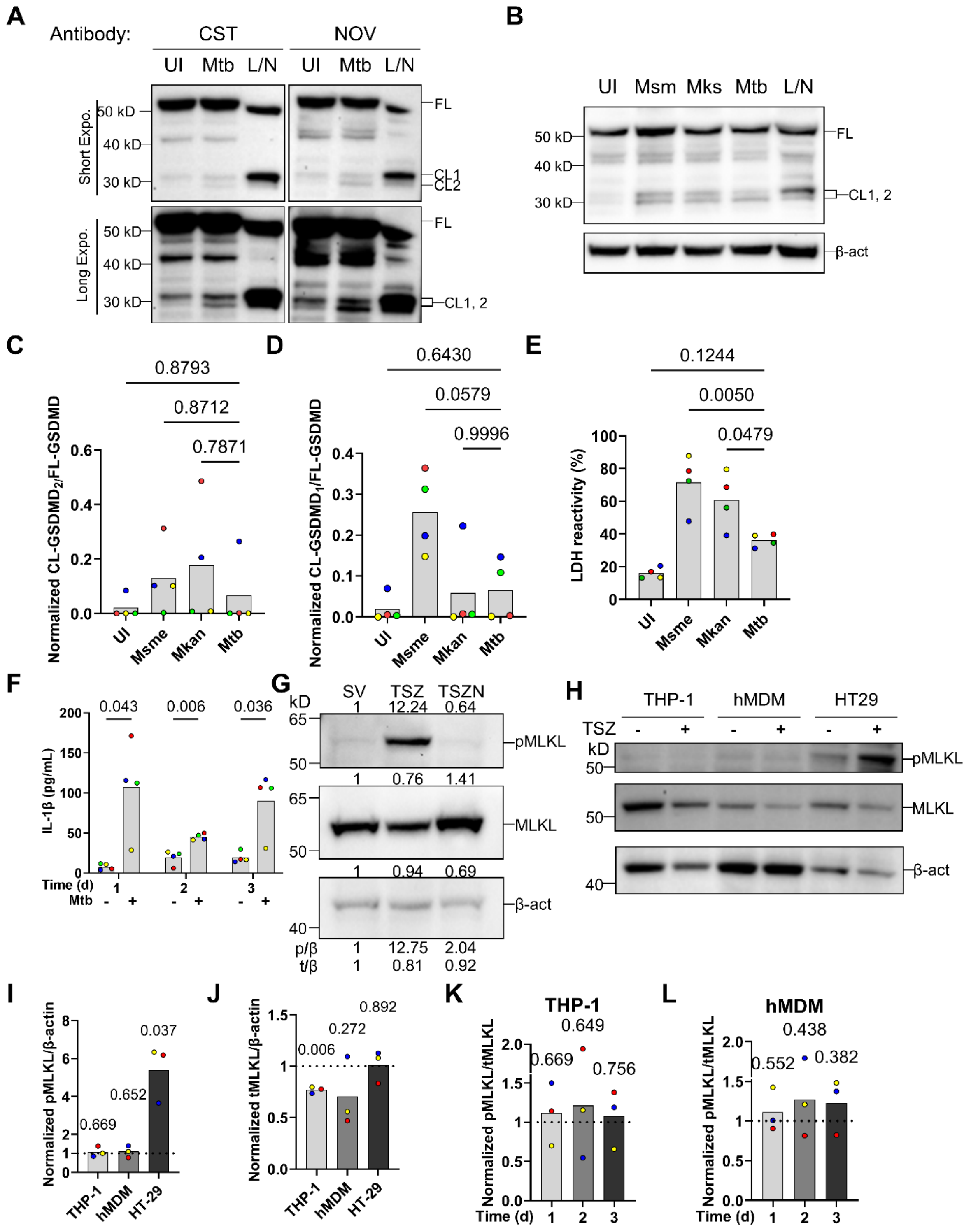

A
Antibody:
CST
NOV
UI Mtb L/N UI Mtb L/N
Short Expo.
Long Expo.
50 kD
40 kD
30 kD
FL
CL1
CL2
CL1, 2
B
UI Msm Mks Mtb L/N
FL
CL1, 2
β-act
C
Normalized CL-GSDMD2/FL-GSDMD
0.8793
0.8712
0.7871
UI Msme Mkan Mtb
D
Normalized CL-GSDMD1/FL-GSDMD
0.6430
0.0579
0.9996
E
LDH reactivity (%)
0.1244
0.0050
0.0479
F
IL-1β (pg/mL)
0.043 0.006 0.036
Time (d)
Mtb
G
kD
SV TSZ TSZN
1 12.24 0.64
pMLKL
1 0.76 1.41
MLKL
1 0.94 0.69
β-act
p/β 1 12.75 2.04
t/β 1 0.81 0.92
H
THP-1 hMDM HT29
TSZ
pMLKL
MLKL
β-act
I
Normalized pMLKL/β-actin
0.669 0.652 0.037
THP-1 hMDM HT-29
J
Normalized tMLKL/β-actin
0.006 0.272 0.892
K
THP-1
Normalized pMLKL/tMLKL
0.669 0.649 0.756
Time (d)
L
hMDM
0.552 0.438 0.382

Fig. S6

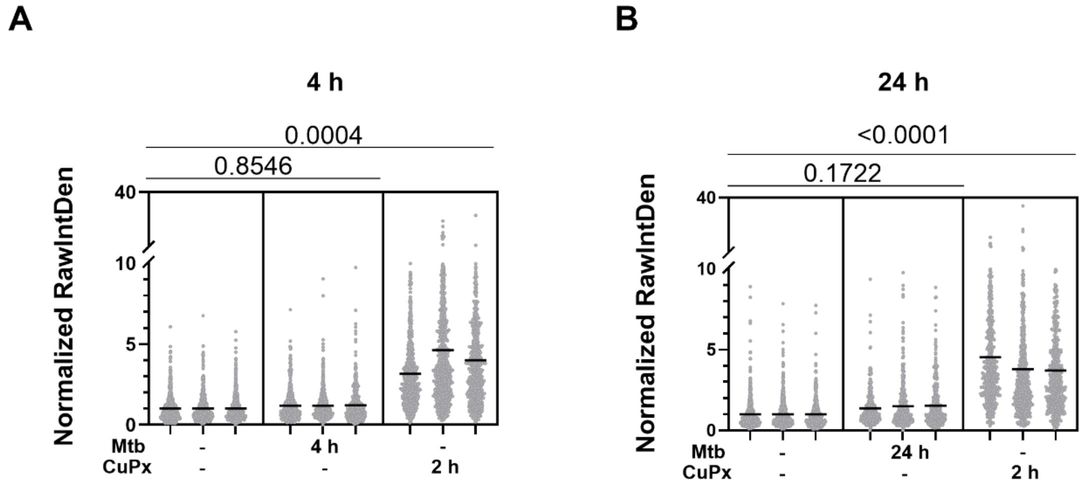
A
4 h
0.0004
0.8546
Normalized RawIntDen
40
10
5
0
Mtb
CuPx
-
-
4 h
-
-
2 h
B
24 h
<0.0001
0.1722
Normalized RawIntDen
40
10
5
0
Mtb
CuPx
-
-
24 h
-
-
2 h

### Fig. S7

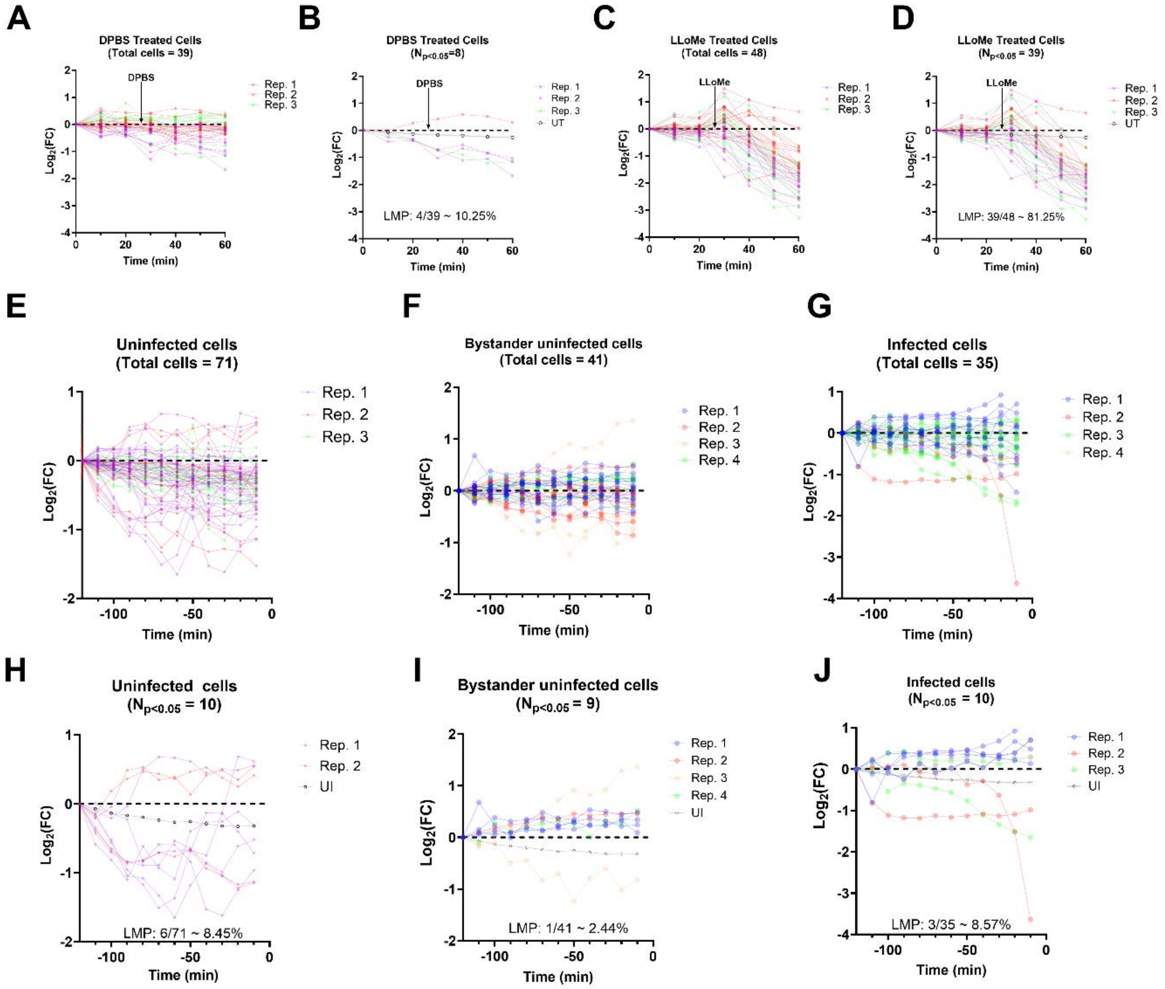
A
DPBS Treated Cells
(Total cells = 39)
B
DPBS Treated Cells
LMP: 4/39 ~ 10.25%
C
LLoMe Treated Cells
(Total cells = 48)
D
LLoMe Treated Cells
LMP: 39/48 ~ 81.25%
E
Uninfected cells
(Total cells = 71)
F
Bystander uninfected cells
(Total cells = 41)
G
Infected cells
(Total cells = 35)
H
Uninfected cells
LMP: 6/71 ~ 8.45%
I
Bystander uninfected cells
LMP: 1/41 ~ 2.44%
J
Infected cells
LMP: 3/35 ~ 8.57%
Time (min)
Rep. 1
Rep. 2
Rep. 3
Rep. 4
UT
UI

Fig. S8

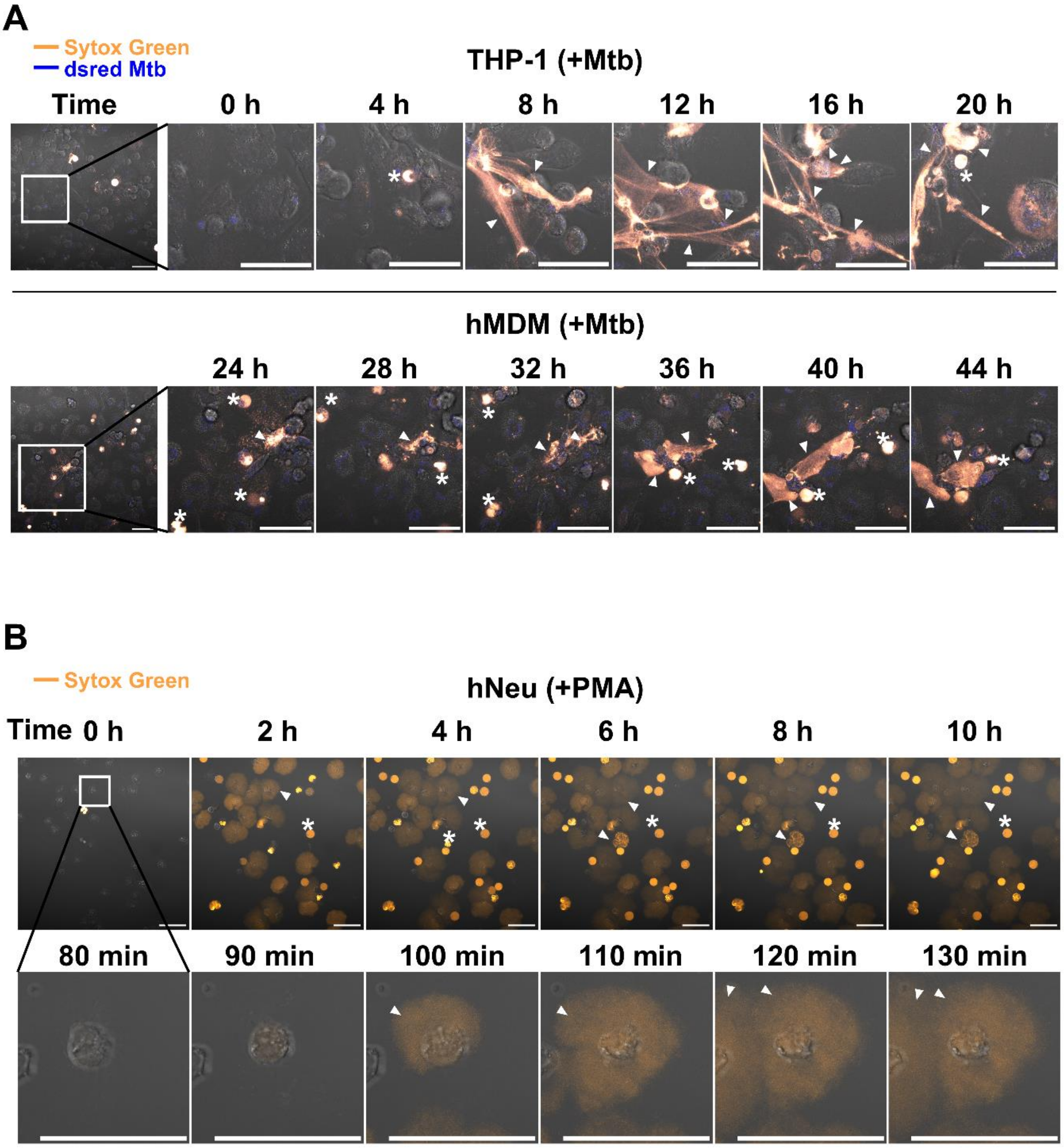

A
Sytox Green
dsred Mtb
THP-1 (+Mtb)
Time
0 h
4 h
8 h
12 h
16 h
20 h
hMDM (+Mtb)
24 h
28 h
32 h
36 h
40 h
44 h
B
Sytox Green
hNeu (+PMA)
Time
0 h
2 h
4 h
6 h
8 h
10 h
80 min
90 min
100 min
110 min
120 min
130 min

Fig. S9

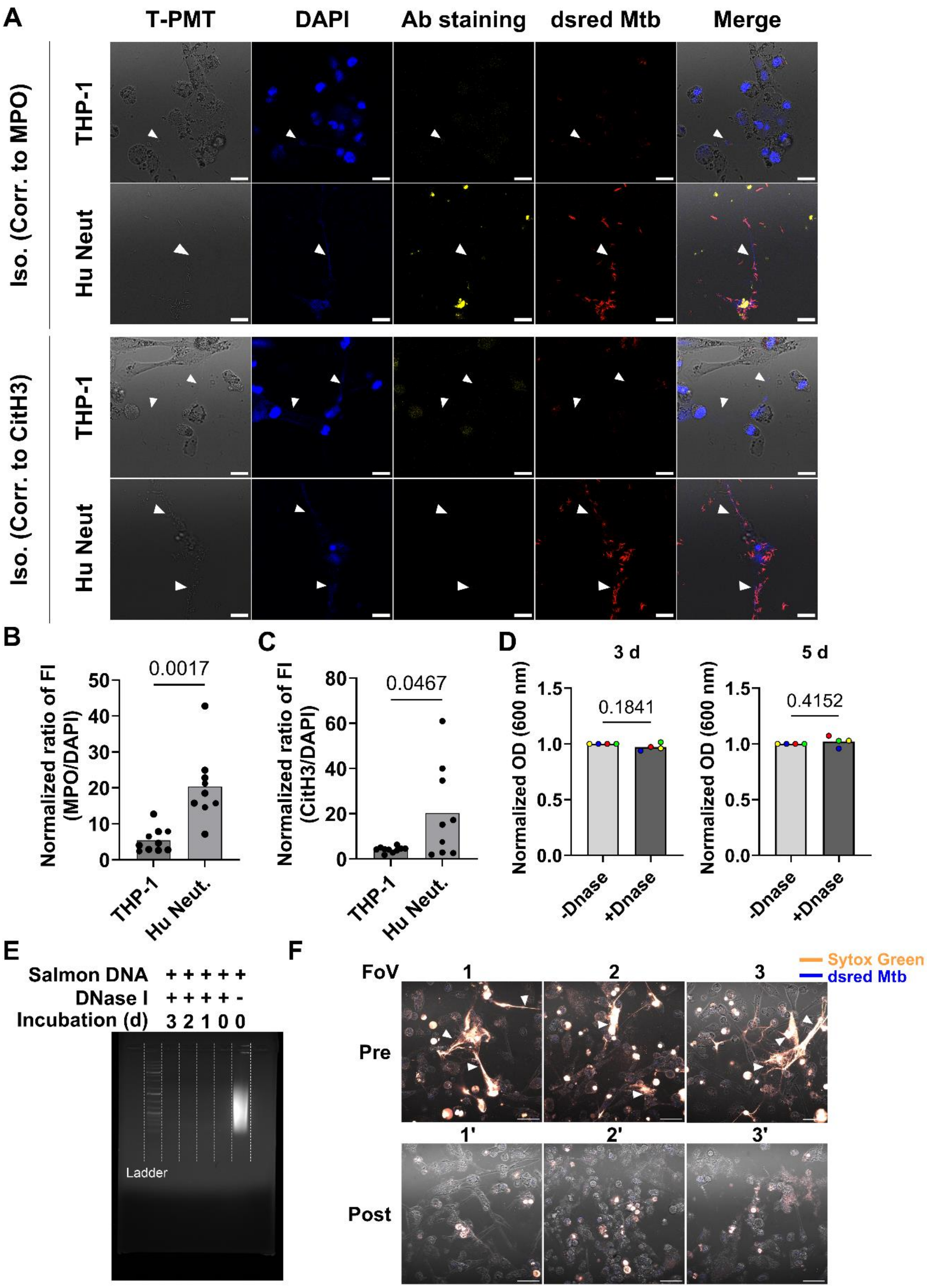

A
T-PMT
DAPI
Ab staining
dsred Mtb
Merge
Iso. (Corr. to MPO)
THP-1
Hu Neut
Iso. (Corr. to CitH3)
THP-1
Hu Neut
B
Normalized ratio of FI (MPO/DAPI)
0.0017
THP-1
Hu Neut.
C
Normalized ratio of FI (CitH3/DAPI)
0.0467
THP-1
Hu Neut.
D
3 d
5 d
Normalized OD (600 nm)
0.1841
0.4152
-Dnase
+Dnase
E
Salmon DNA + + + + + +
DNase I + + + + + -
Incubation (d) 3 2 1 0 0
Ladder
F
Sytox Green
dsred Mtb
FoV
1
2
3
Pre
1'
2'
3'
Post

**Fig. S10**

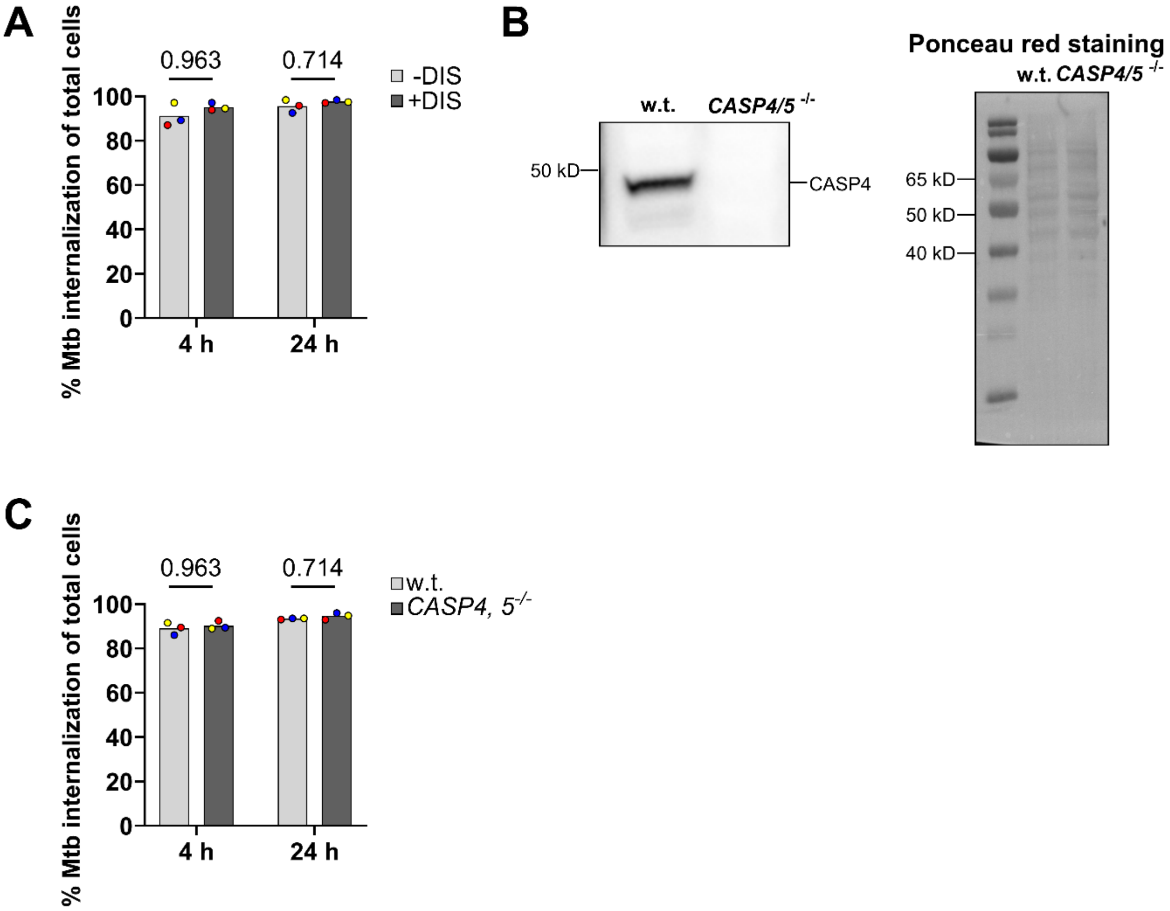
A
% Mtb internalization of total cells
0.963
0.714
-DIS
+DIS
100
80
60
40
20
0
4 h
24 h
B
w.t.
CASP4/5 -/-
50 kD
CASP4
Ponceau red staining
w.t. CASP4/5 -/-
65 kD
50 kD
40 kD
C
% Mtb internalization of total cells
0.963
0.714
w.t.
CASP4, 5-/-
100
80
60
40
20
0
4 h
24 h

### Fig. S11

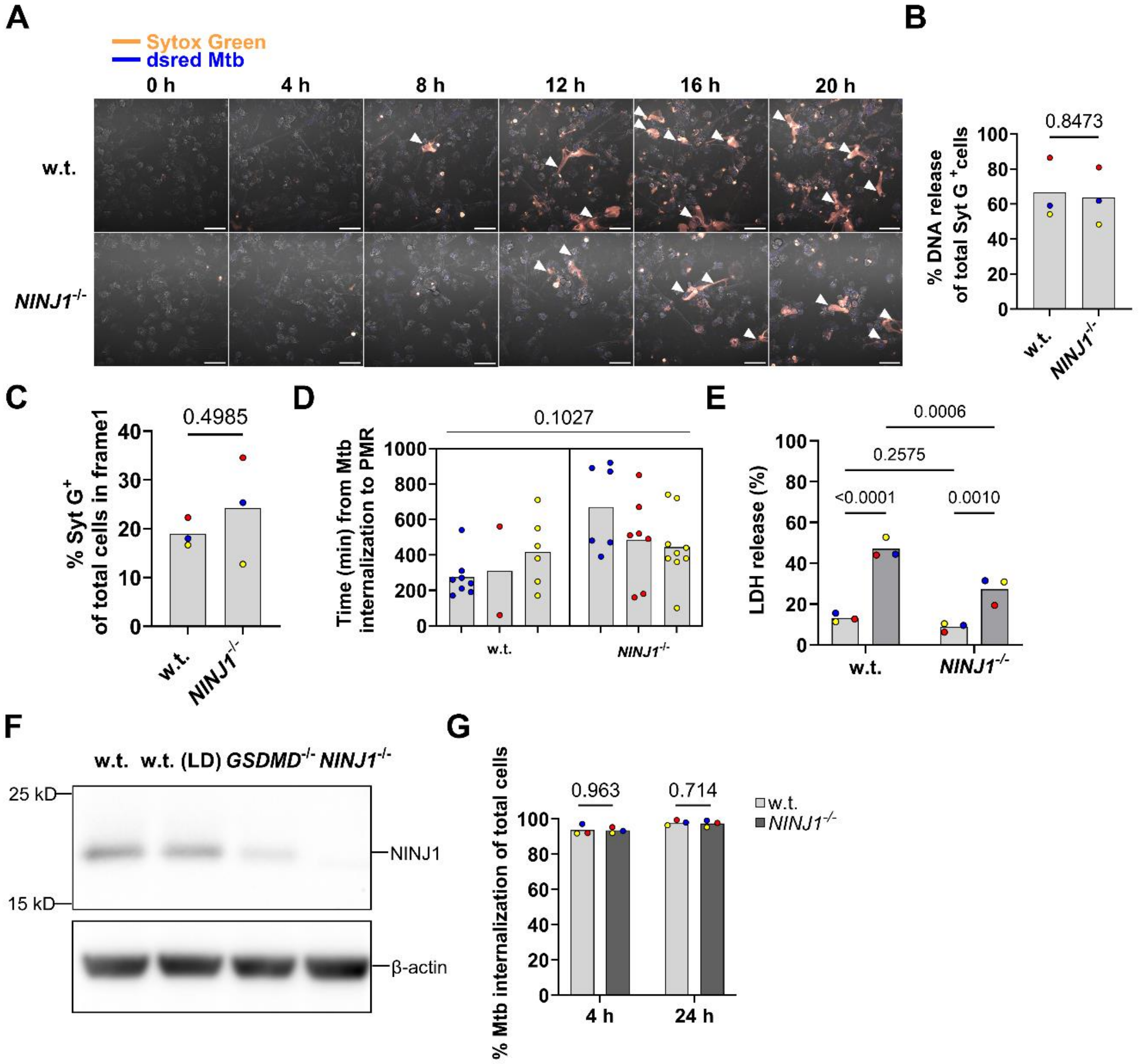
A
Sytox Green
dsred Mtb
0 h
4 h
8 h
12 h
16 h
20 h
w.t.
NINJ1-/-
B
0.8473
% DNA release of total Syt G+ cells
w.t.
NINJ1-/-
C
0.4985
% Syt G+ of total cells in frame1
D
0.1027
Time (min) from Mtb internalization to PMR
E
0.0006
0.2575
<0.0001
0.0010
LDH release (%)
F
w.t.
w.t. (LD)
GSDMD-/-
NINJ1-/-
25 kD
15 kD
NINJ1
β-actin
G
0.963
0.714
w.t.
NINJ1-/-
% Mtb internalization of total cells
4 h
24 h
