## Supplementary material for "Characterization of programmed cell death pathways activated in *Mycobacterium tuberculosis*-infected human macrophages": Uncropped blots.pptx

### Slide 1
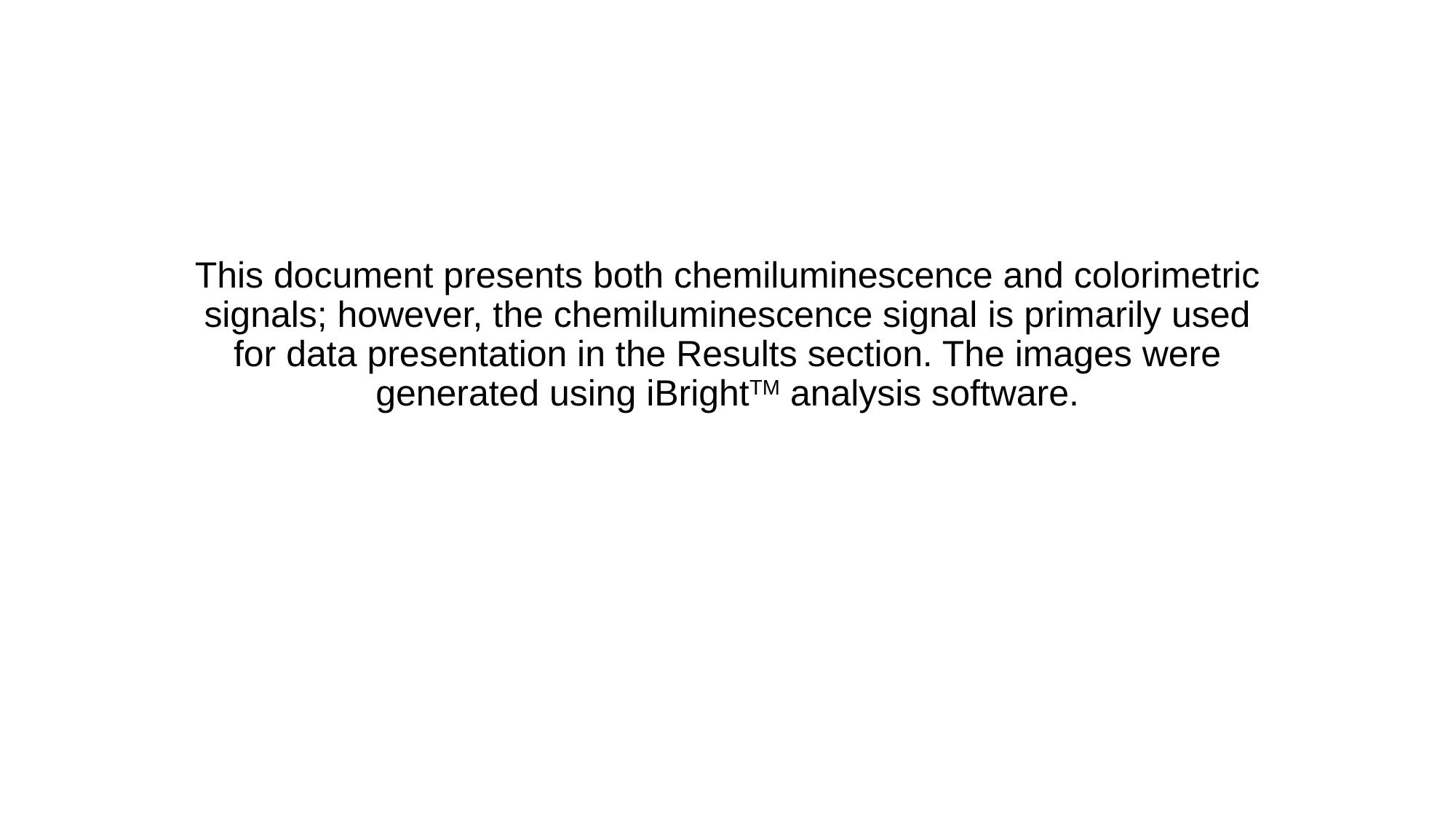

#### This document presents both chemiluminescence and colorimetric signals; however, the chemiluminescence signal is primarily used for data presentation in the Results section. The images were generated using iBright^TM^ analysis software.

### Slide 2
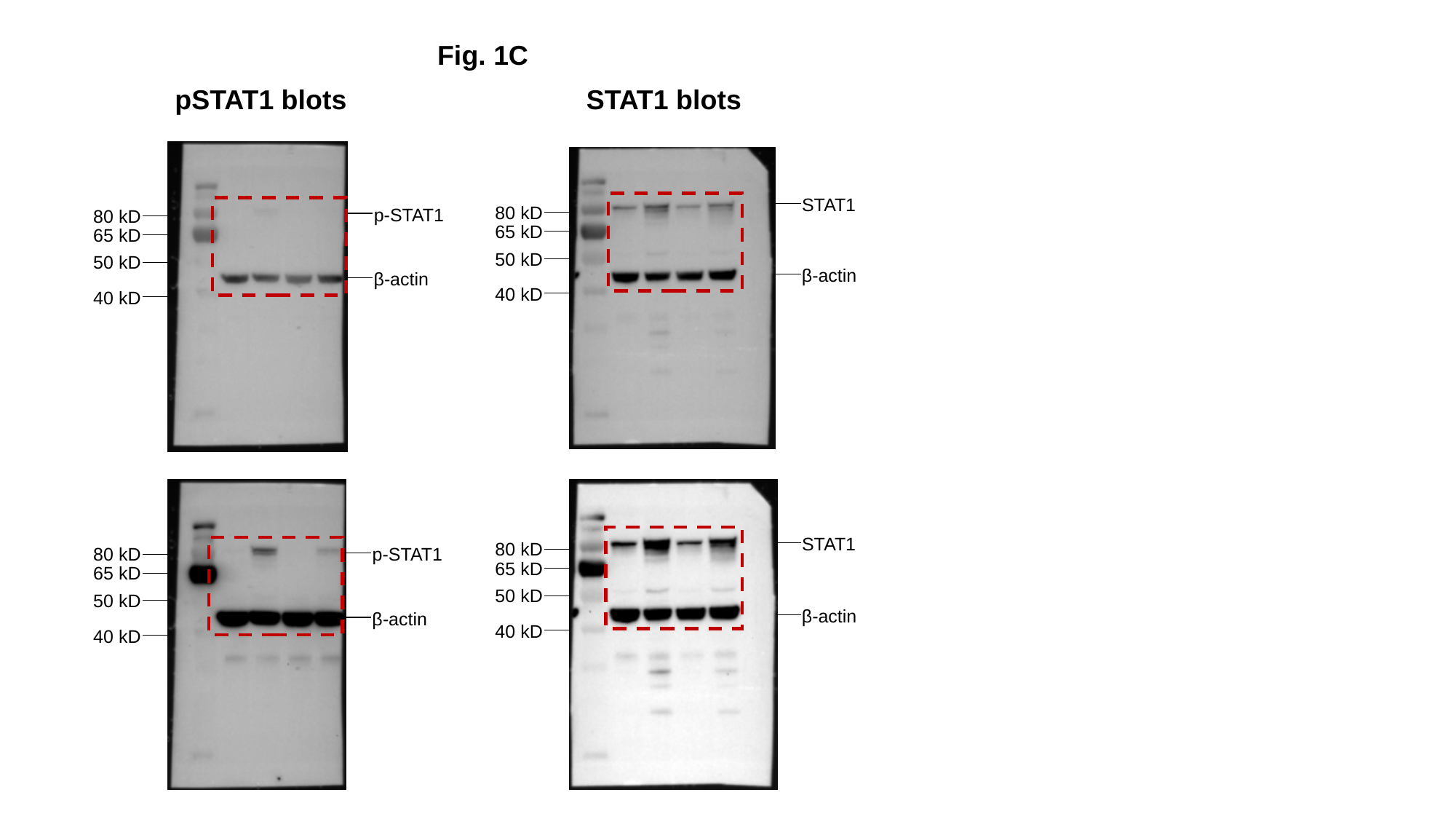

Fig. 1C
pSTAT1 blots
STAT1 blots
STAT1
80 kD
p-STAT1
80 kD
65 kD
65 kD
50 kD
50 kD
β-actin
β-actin
40 kD
40 kD
STAT1
80 kD
p-STAT1
80 kD
65 kD
65 kD
50 kD
50 kD
β-actin
β-actin
40 kD
40 kD

### Slide 3
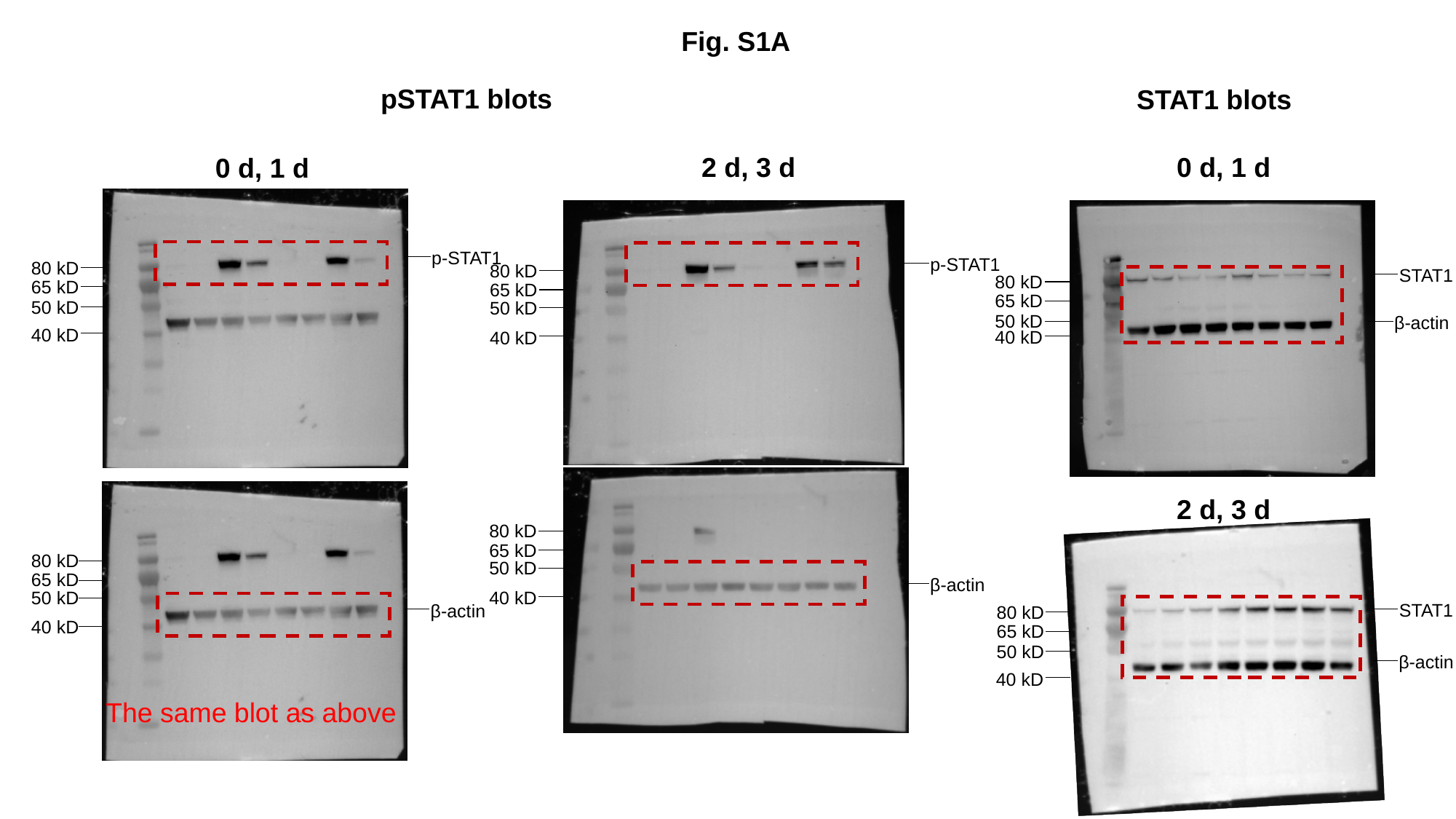

Fig. S1A
pSTAT1 blots
STAT1 blots
2 d, 3 d
0 d, 1 d
0 d, 1 d
p-STAT1
p-STAT1
80 kD
80 kD
STAT1
80 kD
65 kD
65 kD
65 kD
50 kD
50 kD
50 kD
β-actin
40 kD
40 kD
40 kD
2 d, 3 d
80 kD
65 kD
80 kD
50 kD
65 kD
β-actin
50 kD
40 kD
STAT1
β-actin
80 kD
40 kD
65 kD
50 kD
β-actin
40 kD
The same blot as above

### Slide 4
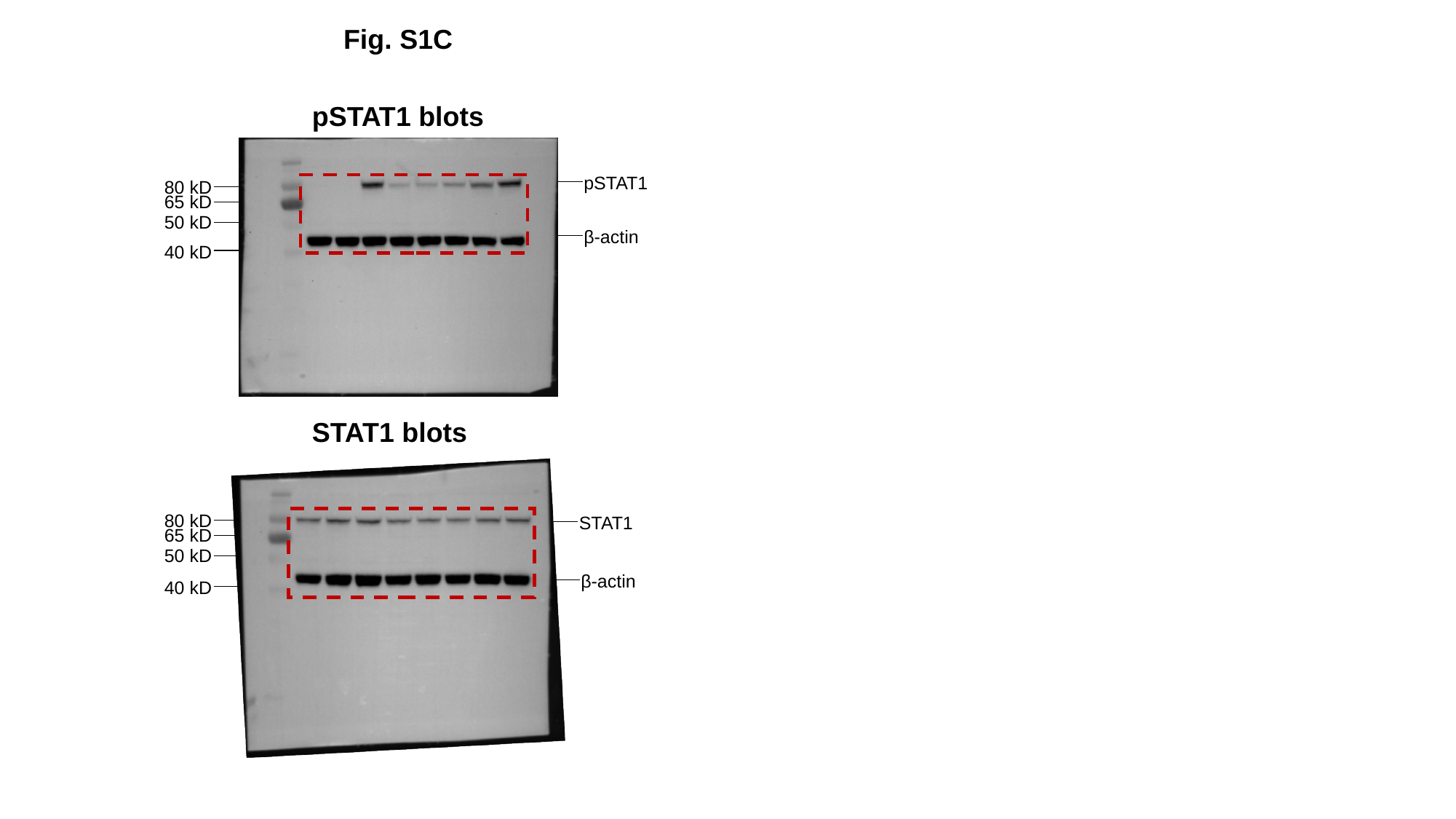

Fig. S1C
pSTAT1 blots
pSTAT1
80 kD
65 kD
50 kD
β-actin
40 kD
STAT1 blots
80 kD
STAT1
65 kD
50 kD
β-actin
40 kD

### Slide 5
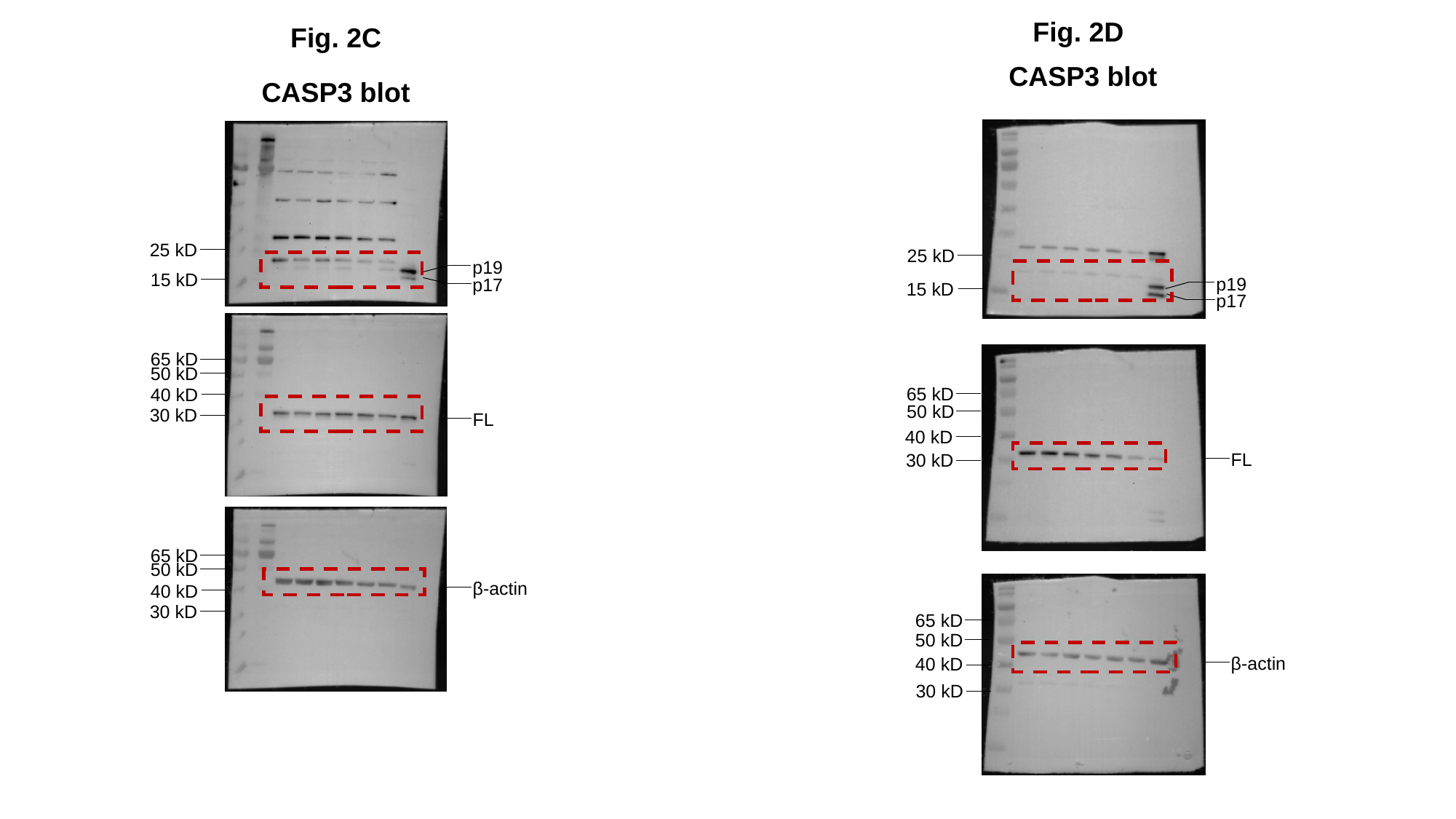

Fig. 2D
Fig. 2C
CASP3 blot
CASP3 blot
25 kD
25 kD
p19
15 kD
p19
p17
15 kD
p17
65 kD
50 kD
65 kD
40 kD
50 kD
30 kD
FL
40 kD
FL
30 kD
65 kD
50 kD
β-actin
40 kD
30 kD
65 kD
50 kD
β-actin
40 kD
30 kD

### Slide 6
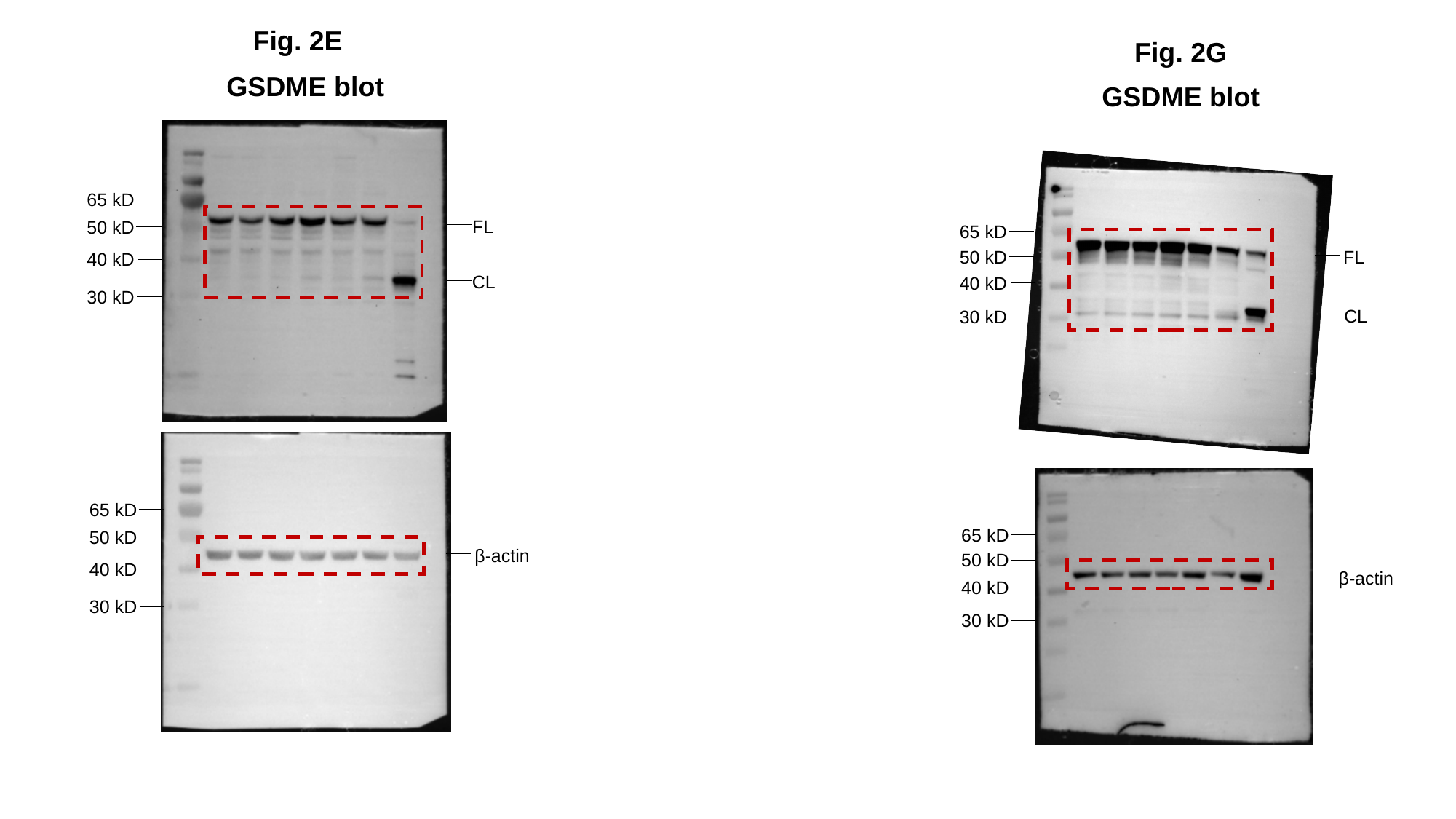

Fig. 2E
Fig. 2G
GSDME blot
GSDME blot
65 kD
FL
50 kD
65 kD
50 kD
FL
40 kD
CL
40 kD
30 kD
CL
30 kD
65 kD
65 kD
50 kD
β-actin
50 kD
40 kD
β-actin
40 kD
30 kD
30 kD

### Slide 7
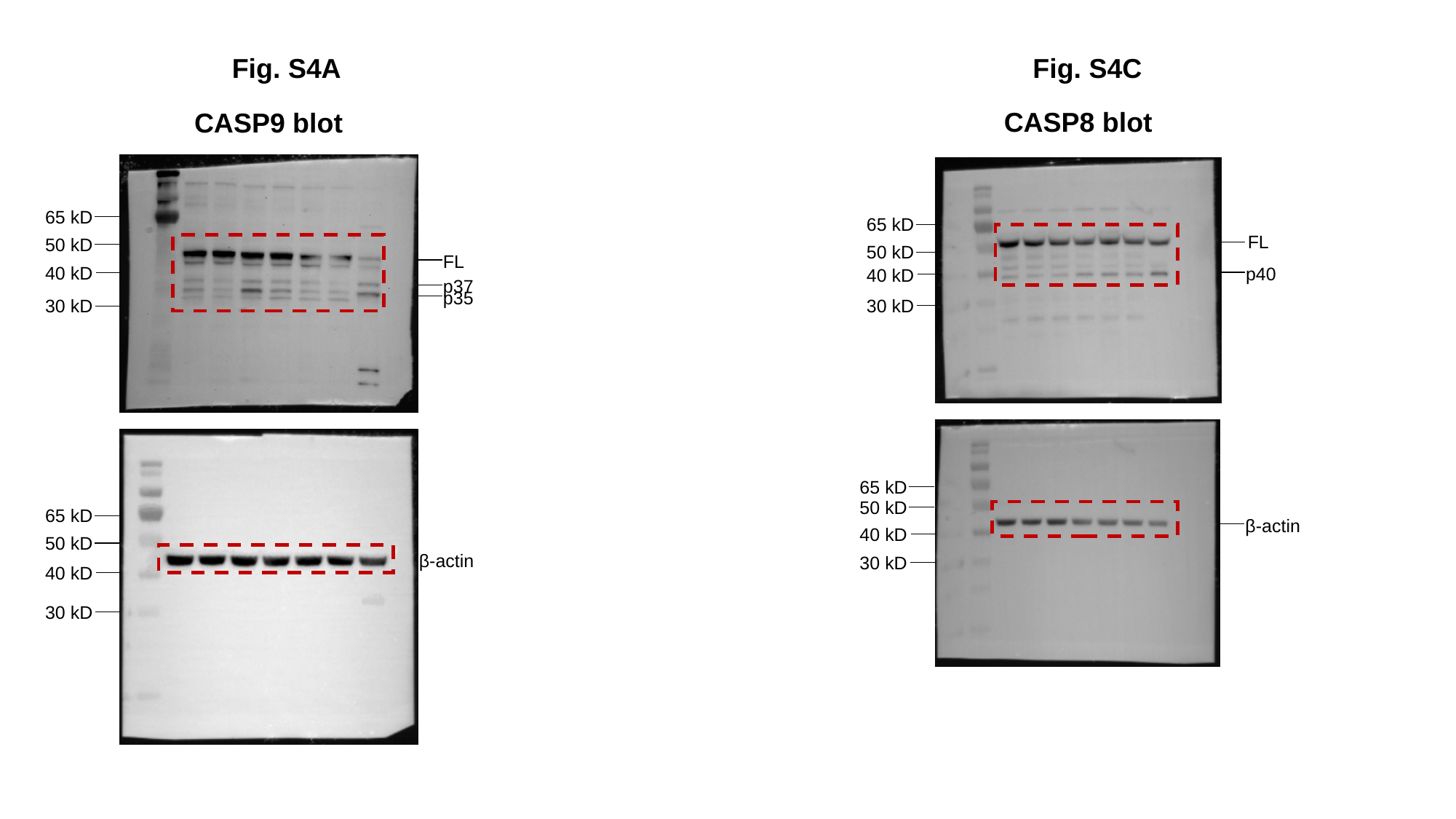

Fig. S4C
Fig. S4A
CASP8 blot
CASP9 blot
65 kD
65 kD
FL
50 kD
50 kD
FL
40 kD
p40
40 kD
p37
p35
30 kD
30 kD
65 kD
50 kD
65 kD
β-actin
40 kD
50 kD
β-actin
30 kD
40 kD
30 kD

### Slide 8
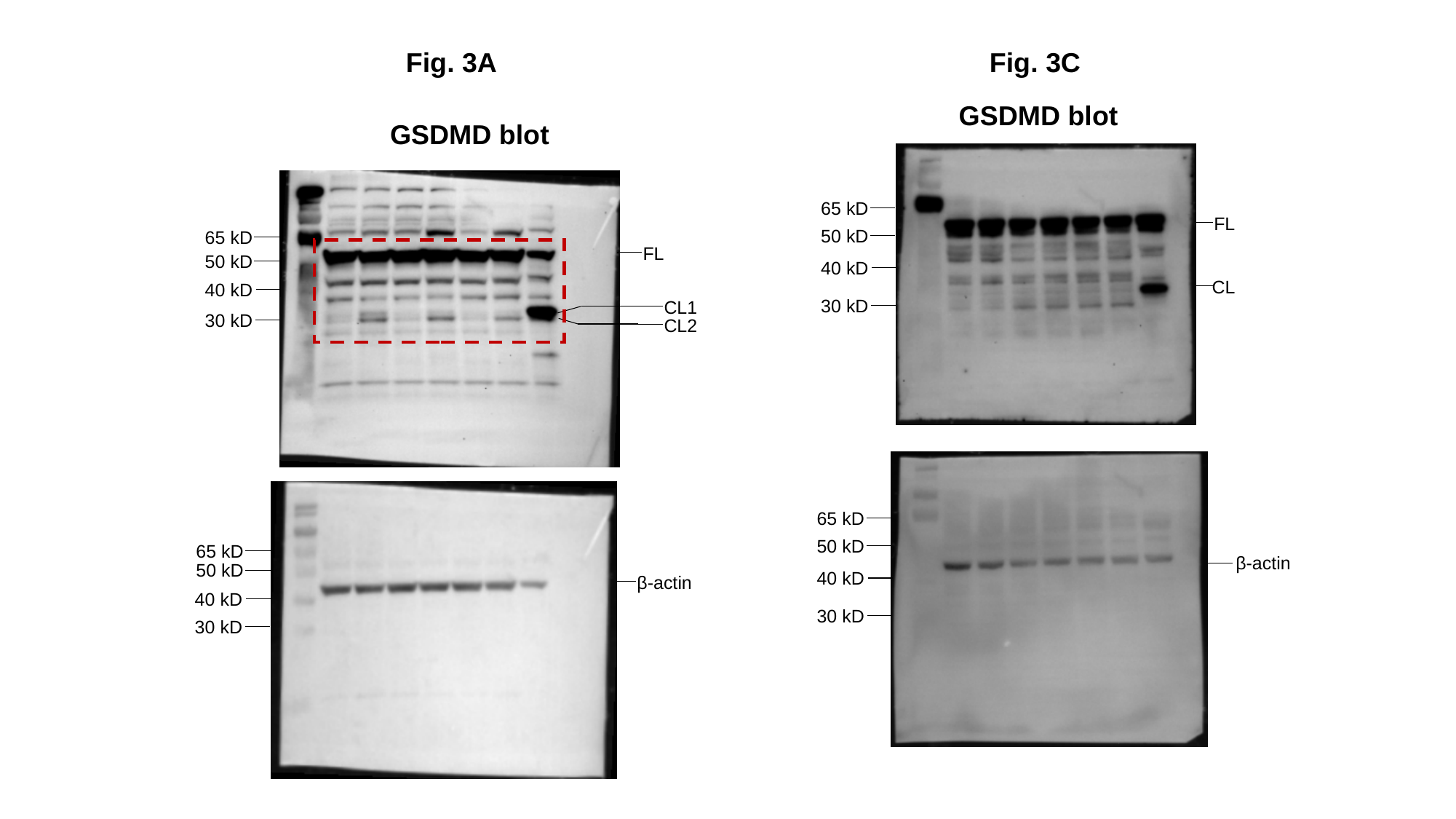

Fig. 3A
Fig. 3C
GSDMD blot
GSDMD blot
65 kD
FL
50 kD
65 kD
FL
50 kD
40 kD
CL
40 kD
30 kD
CL1
30 kD
CL2
65 kD
50 kD
65 kD
β-actin
50 kD
40 kD
β-actin
40 kD
30 kD
30 kD

### Slide 9
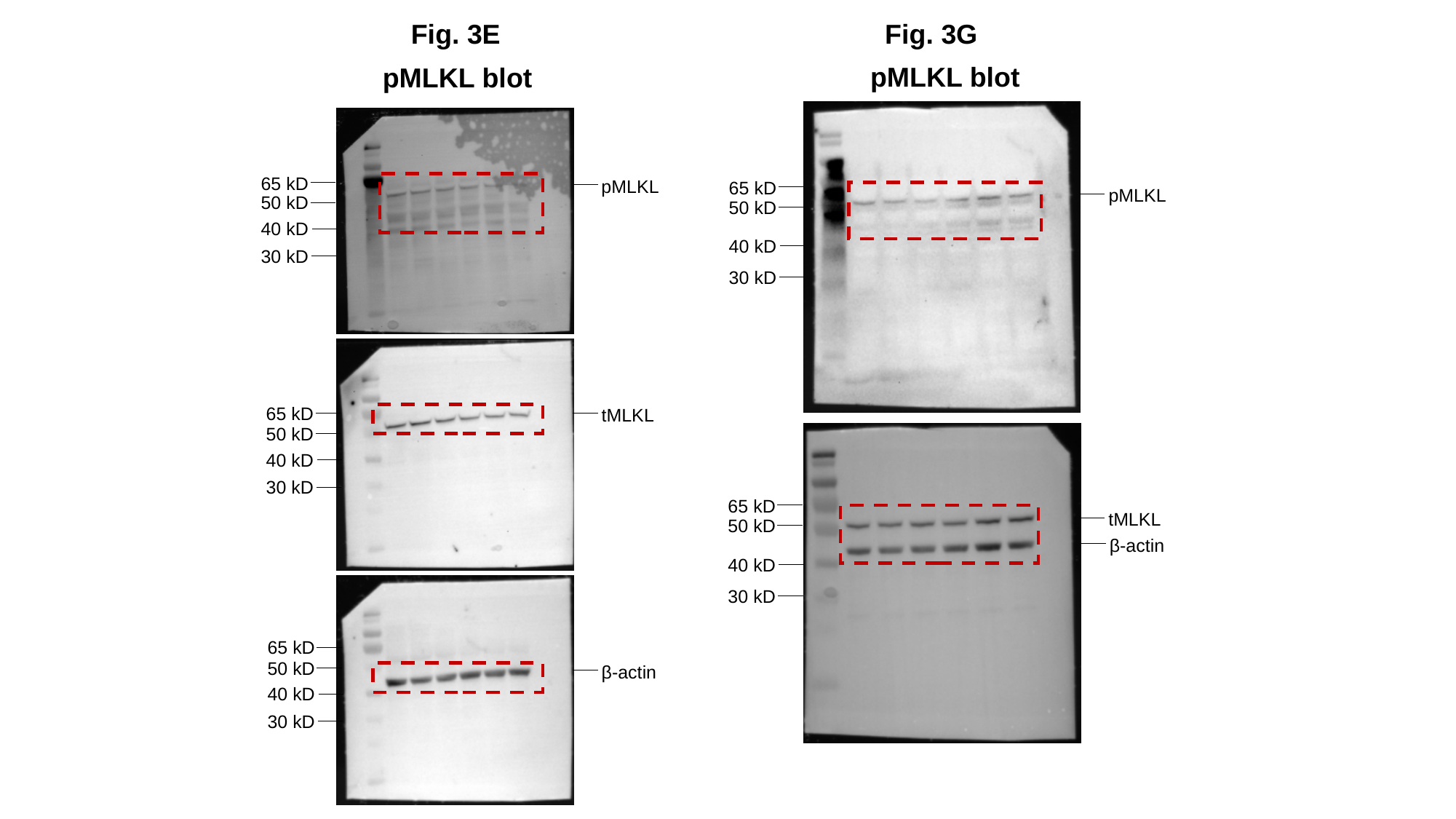

Fig. 3E
Fig. 3G
pMLKL blot
pMLKL blot
65 kD
pMLKL
65 kD
pMLKL
50 kD
50 kD
40 kD
40 kD
30 kD
30 kD
65 kD
tMLKL
50 kD
40 kD
30 kD
65 kD
tMLKL
50 kD
β-actin
40 kD
30 kD
65 kD
50 kD
β-actin
40 kD
30 kD

### Slide 10
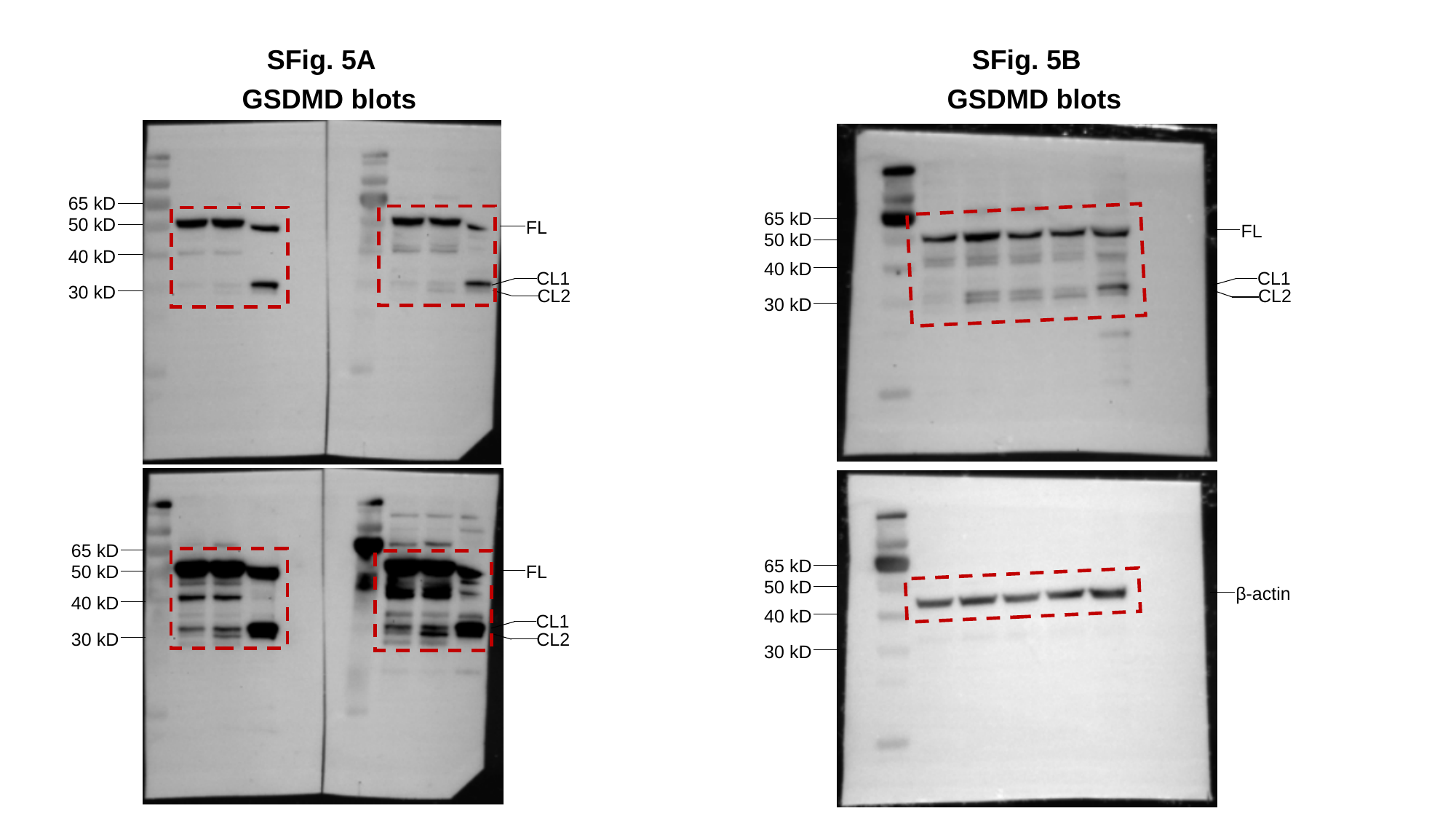

SFig. 5A
SFig. 5B
GSDMD blots
GSDMD blots
65 kD
65 kD
50 kD
FL
FL
50 kD
40 kD
40 kD
CL1
CL1
30 kD
CL2
CL2
30 kD
65 kD
65 kD
50 kD
FL
50 kD
β-actin
40 kD
40 kD
CL1
30 kD
CL2
30 kD

### Slide 11
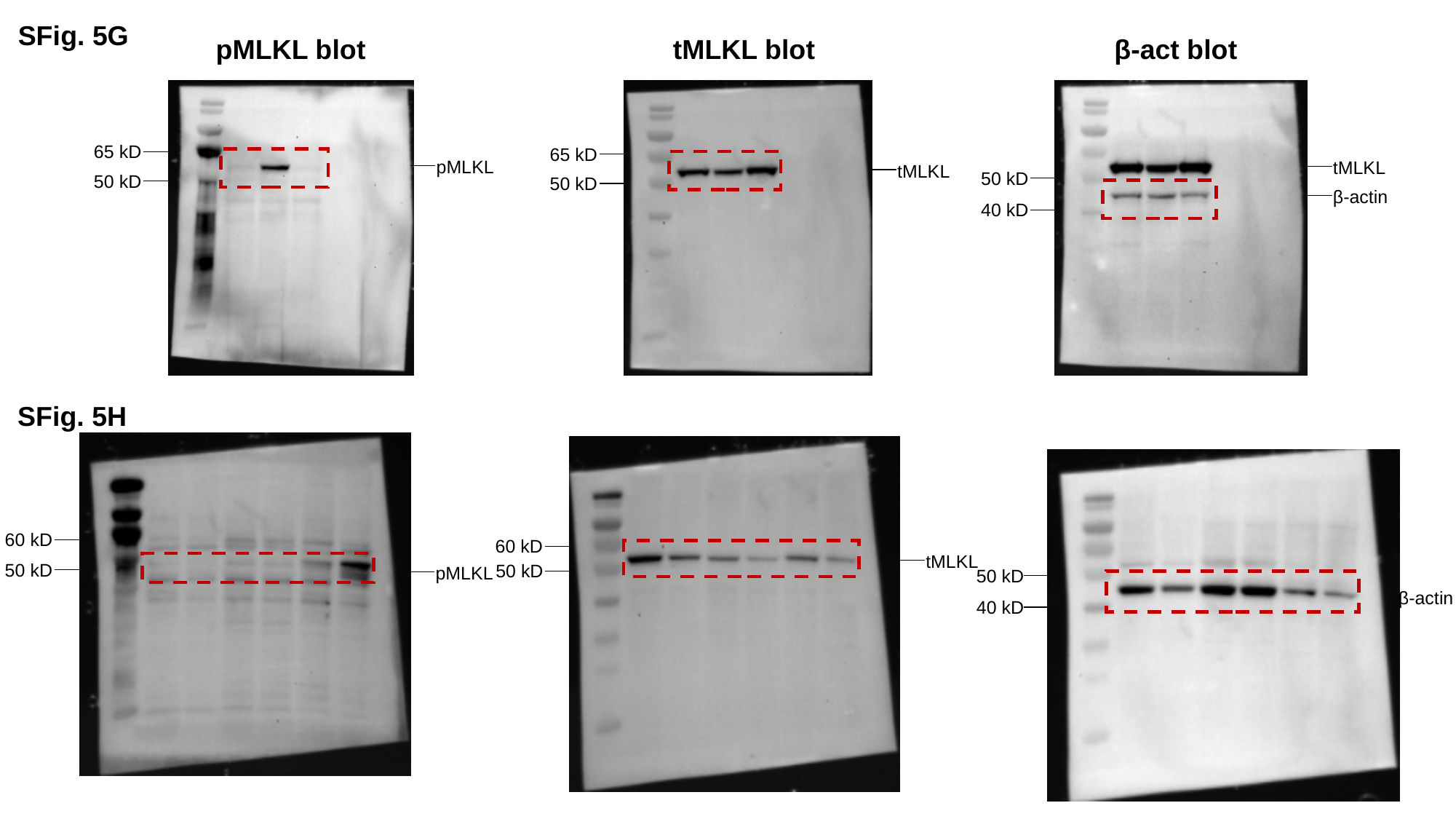

SFig. 5G
pMLKL blot
tMLKL blot
β-act blot
65 kD
65 kD
pMLKL
tMLKL
tMLKL
50 kD
50 kD
50 kD
β-actin
40 kD
SFig. 5H
60 kD
60 kD
tMLKL
50 kD
50 kD
pMLKL
50 kD
β-actin
40 kD

### Slide 12
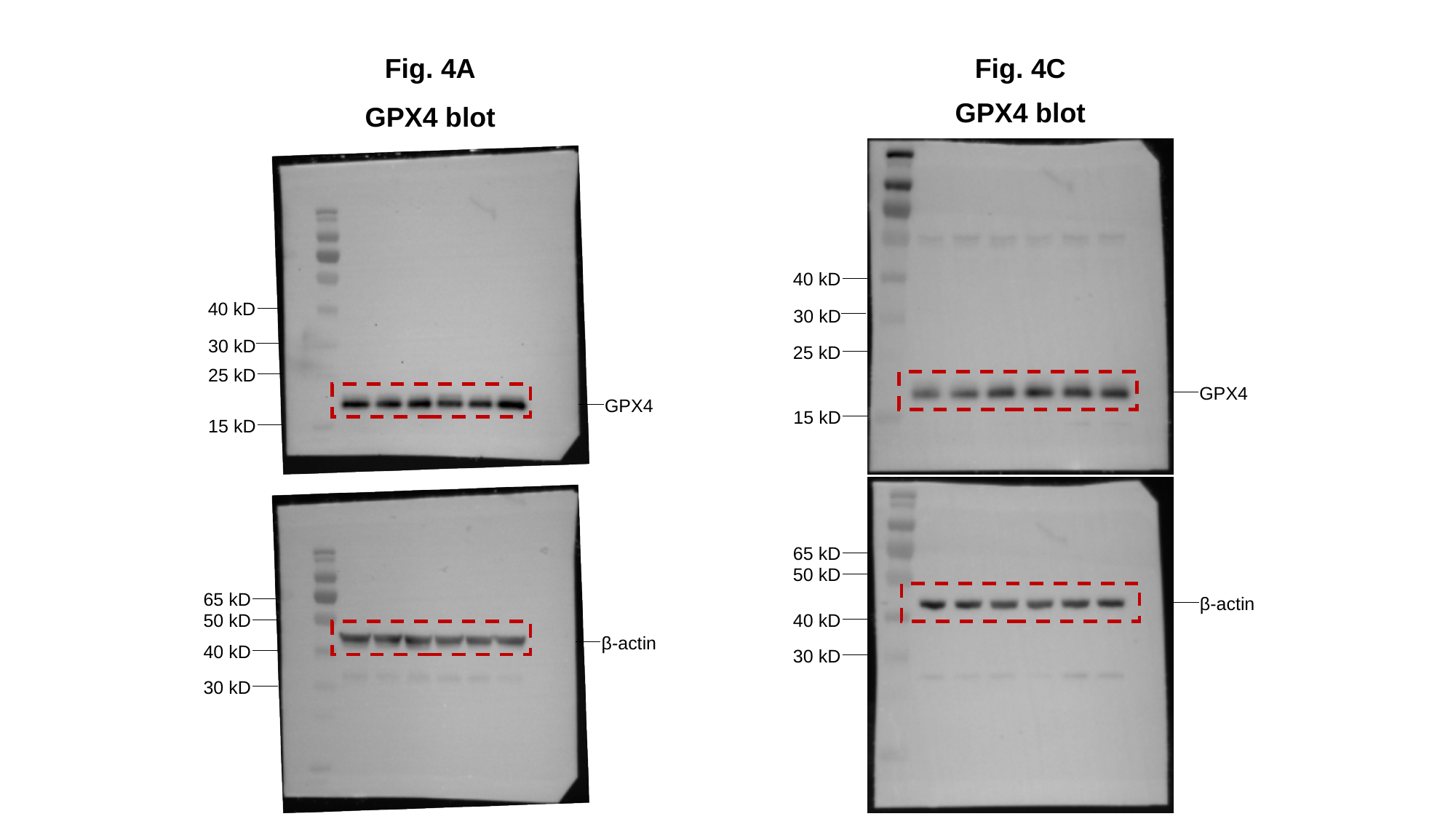

Fig. 4A
Fig. 4C
GPX4 blot
GPX4 blot
40 kD
40 kD
30 kD
30 kD
25 kD
25 kD
GPX4
GPX4
15 kD
15 kD
65 kD
50 kD
65 kD
β-actin
50 kD
40 kD
β-actin
40 kD
30 kD
30 kD

### Slide 13
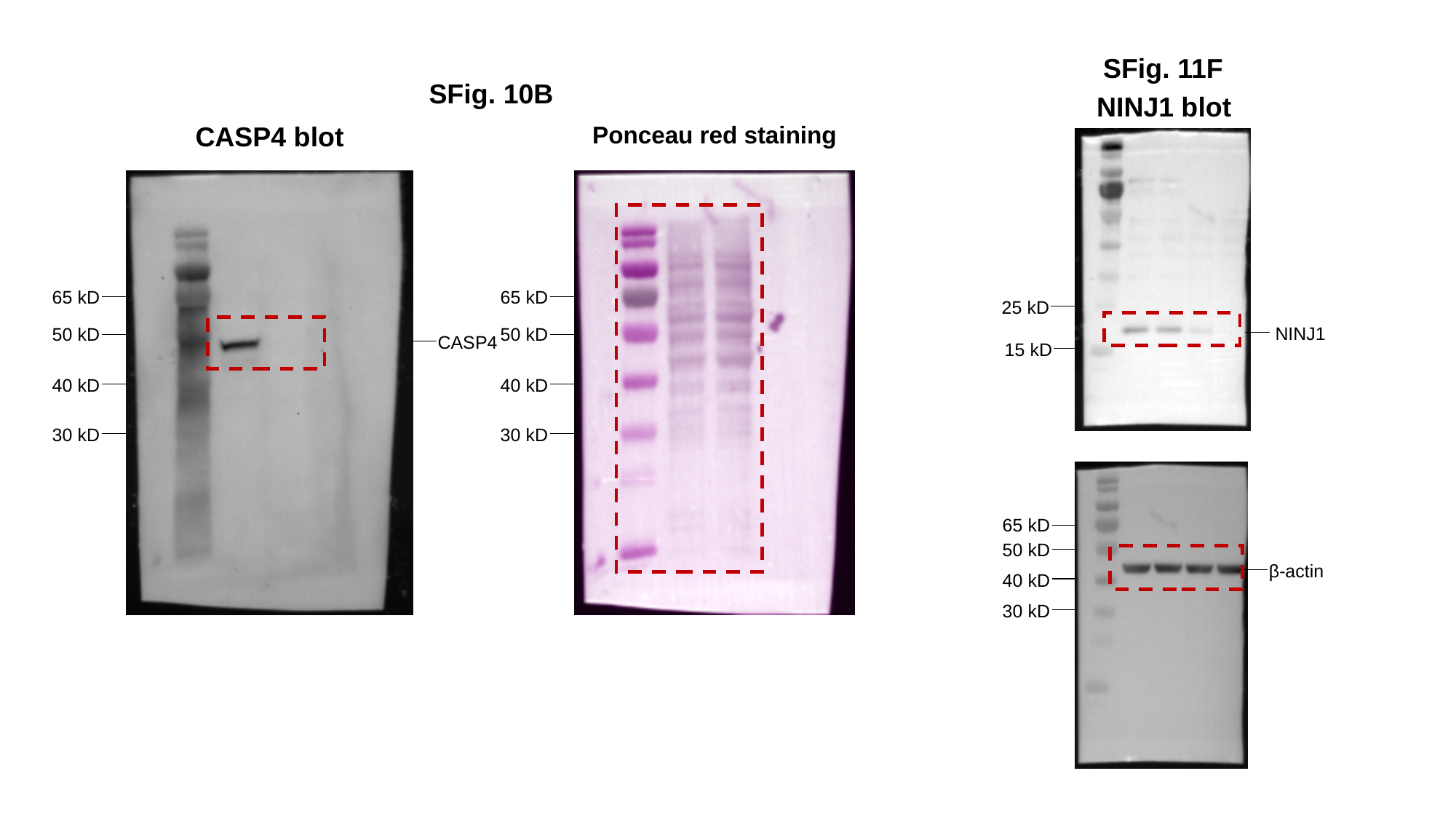

SFig. 11F
SFig. 10B
NINJ1 blot
CASP4 blot
Ponceau red staining
65 kD
65 kD
25 kD
NINJ1
50 kD
50 kD
CASP4
15 kD
40 kD
40 kD
30 kD
30 kD
65 kD
50 kD
β-actin
40 kD
30 kD
